# Integrating protein networks and machine learning for disease stratification in the Hereditary Spastic Paraplegias

**DOI:** 10.1101/2021.01.14.425874

**Authors:** Nikoleta Vavouraki, James E. Tomkins, Eleanna Kara, Henry Houlden, John Hardy, Marcus J. Tindall, Patrick A. Lewis, Claudia Manzoni

**Affiliations:** Department of Pharmacy, University of Reading, Reading, RG6 6AH, United Kingdom; Department of Mathematics and Statistics, University of Reading, Reading, RG6 6AH, United Kingdom; Department of Neuromuscular Diseases, UCL Queen Square Institute of Neurology, London, WC1N 3BG, United Kingdom; Department of Neurodegenerative Disease, UCL Queen Square Institute of Neurology, London, WC1N 3BG, United Kingdom; Institute of Cardiovascular and Metabolic Research, University of Reading, Reading, RG6 6AS, United Kingdom; Department of Comparative Biomedical Sciences, Royal Veterinary College, London, NW1 0TU, United Kingdom; School of Pharmacy, University College London, London, WC1N 1AX, United Kingdom

**Keywords:** Hereditary Spastic Paraplegias, machine learning, neurodegeneration, PINOT, protein interaction network

## Abstract

The Hereditary Spastic Paraplegias are a group of neurodegenerative diseases characterized by spasticity and weakness in the lower body. Despite the identification of causative mutations in over 70 genes, the molecular aetiology remains unclear. Due to the combination of genetic diversity and variable clinical presentation, the Hereditary Spastic Paraplegias are a strong candidate for protein-protein interaction network analysis as a tool to understand disease mechanism(s) and to aid functional stratification of phenotypes. In this study, experimentally validated human protein-protein interactions were used to create a protein-protein interaction network based on the causative Hereditary Spastic Paraplegia genes. Network evaluation as a combination of both topological analysis and functional annotation led to the identification of core proteins in putative shared biological processes such as intracellular transport and vesicle trafficking. The application of machine learning techniques suggested a functional dichotomy linked with distinct sets of clinical presentations, suggesting there is scope to further classify conditions currently described under the same umbrella term of Hereditary Spastic Paraplegias based on specific molecular mechanisms of disease.

## Introduction

The Hereditary Spastic Paraplegias (HSPs) are a group of heterogeneous neurodegenerative diseases characterised by the core features of slowly progressive bilateral lower limb spasticity, hyperreflexia and extensor plantar responses (Harding, 1983) accompanied by degeneration of the upper-motor neurons (Deluca *et al*, 2004). Although the first description of clinical presentations we now refer to as HSPs dates back at least 140 years (Lorrain, 1898; Strümpell, 1880), the molecular mechanisms responsible for disease onset are, to date, still unclear. A number of mechanisms have been proposed to contribute to the degenerative process, including dysfunction of intracellular active transport and endolysosomal trafficking, alteration of lipid metabolism and endoplasmic reticulum shaping as well as disruption of mitochondria homeostasis (Blackstone, 2012, 2018a; Blackstone *et al*, 2011; Boutry *et al*, 2019).

The heterogeneity of the HSPs derives from both the complex range of clinical presentations (summarised in Supplementary Table 1) and diverse underlying genetic causes. Regarding the former, the age of onset can vary from early childhood to late adulthood, all modes of inheritance can be observed, and the form of the disease can be pure or complex. Complex forms of the HSPs are defined by the co-occurrence of clinical features in addition to lower limb spasticity, including peripheral neuropathy, seizures, cognitive impairment and optic atrophy (Fink, 2013). Regarding the genetic heterogeneity of HSPs, mutations in over 70 genes have been associated with the HSPs (Faber *et al*, 2017), rendering it one of the hereditary disorders with the highest numbers of known causative genes (Blackstone, 2018a). In such a complex scenario, it is not clear as to whether all the HSP syndromes, despite being classified under the same umbrella term, share the same underlying molecular aetiology (Blackstone, 2018a). Given the lack of treatments able to prevent, halt or revert the HSPs, understanding the aetiology of these disorders and gaining greater clarity in this area of HSP biology is crucial.

The intersection of genetics and functional biology has, historically, been dominated by single gene investigations, focusing on understanding the role of individual genes in cellular processes and phenotypes. This approach is powerful, but it allows for studying a limited number of genes at a time (Manzoni *et al*, 2020). In contrast, systems biology approaches such as protein-protein interaction (PPI) network (PPIN) analyses provide tools to evaluate the entirety of known genes/proteins involved in a disease collectively through a holistic approach (Koh *et al*, 2012). The connections within the PPIN can be subjected to mathematical analysis to gain insight into the global relationships among potential contributors to the disease process, thus creating an *in silico* model system to investigate the molecular mechanisms and generate hypotheses to further support functional research and disease modelling (Manzoni *et al*., 2020).

This paper describes the first study in which PPINs are created solely based on experimentally validated human PPIs of HSP genes, and are applied to the investigation of HSP pathogenesis to identify global mechanisms, as well as individual processes involved in subtypes of disease following stratification based on the association of specific HSP genes with particular clinical features. Based on a combination of network, functional, and machine learning analyses, we propose HSPs to be subdivided into at least 2 major aetiological groups. These results might suggest that not all the HSPs’ clinical manifestations relate to the same disruption at a molecular level, and that it is indeed possible to hypothesise stratification of HSP patients based on the molecular aspects of disease. This is an *in silico* modelling approach, thus it would require further functional validation; nevertheless it suggests that both drug discovery and clinical trials for HSPs would need to take into consideration the molecular heterogeneity of disease.

## Results

### Generation of PPI networks

The HSP seeds (HSP genes, n=66 and test seeds, n=17; see Table 1, Supplementary Table 1, and Materials and Methods) were used as the input list to query the online tool, PINOT (Tomkins *et al*, 2020), generating a list of experimentally validated, human PPIs. Briefly, PINOT collects PPIs from 7 manually curated databases that fully or partially comply with the IMEx annotation guidelines (Orchard *et al*, 2012), and scores each interaction based on the number of different methods and publications in which it has been described. PPIs with a final PINOT score <3 were excluded from further analyses as these interactions lack replication in the curated literature (i.e. they are reported in only 1 publication and detected by only 1 method). Following this filter, 746 interactors of HSP seeds were retained. Of note, 15 of the initial seeds were excluded due to no PPIs being identified (a total of 57 HSP seeds and 11 test seeds were retained). The resulted filtered network was termed the global HSP-PPIN and was composed of 814 nodes (57 HSP seeds + 11 test seeds + 746 interactors) connected *via* 925 edges (Supplementary File 1). The global HSP-PPIN (Supplementary Fig 1) was composed of 1 main graph that contained the majority of nodes (n=755/814, 92.8%), including the majority of seeds (n=53/68, 77.9%) and 14 additional unconnected, smaller graphs. Of particular note is the presence of an interactor in the global HSP-PPIN, RNF170, which was found to be associated with the HSPs (i.e. an additional HSP gene) in a study published after the creation of the network (Wagner *et al*, 2019).

**Table 1.**
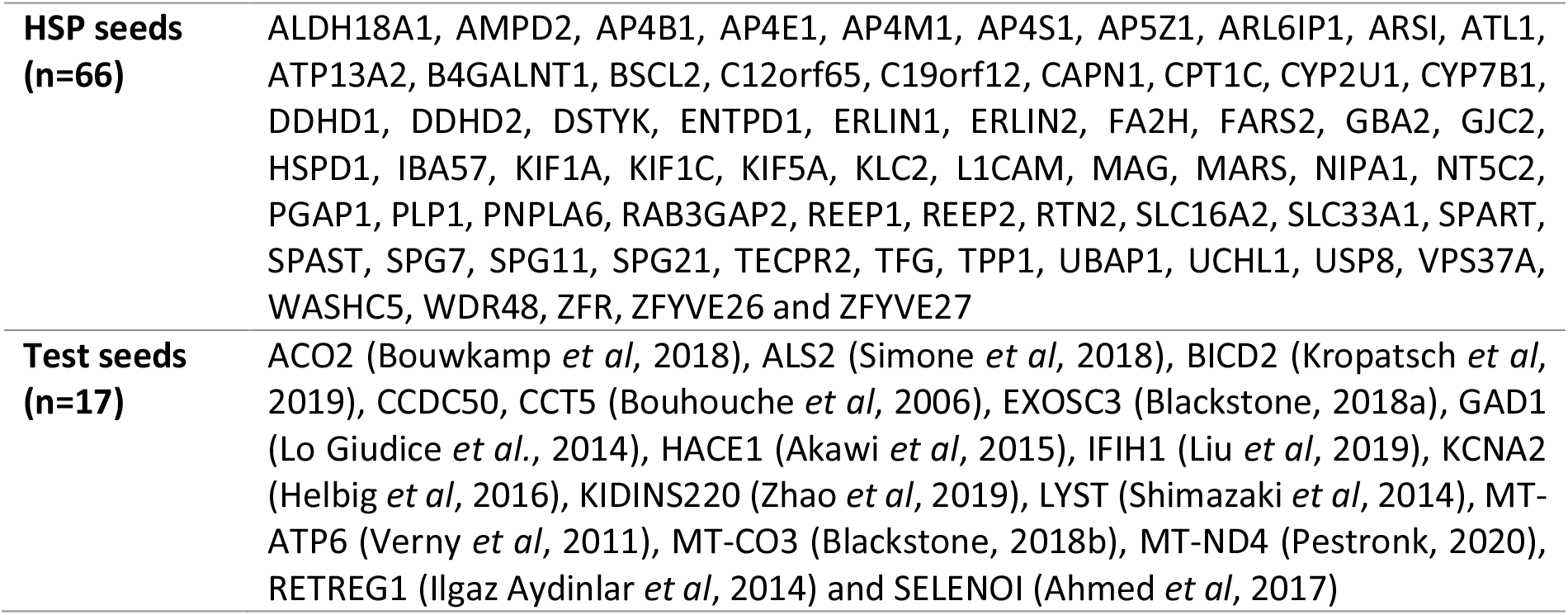
Seeds for the HSP network.

|  |  |
| --- | --- |
| **HSP seeds**<br>**(n=66)** | ALDH18A1, AMPD2, AP4B1, AP4E1, AP4M1, AP4S1, AP5Z1, ARL6IP1, ARSI, ATL1, ATP13A2, B4GALNT1, BSCL2, C12orf65, C19orf12, CAPN1, CPT1C, CYP2U1, CYP7B1, DDHD1, DDHD2, DSTYK, ENTPD1, ERLIN1, ERLIN2, FA2H, FARS2, GBA2, GJC2, HSPD1, IBA57, KIF1A, KIF1C, KIF5A, KLC2, L1CAM, MAG, MARS, NIPA1, NT5C2, PGAP1, PLP1, PNPLA6, RAB3GAP2, REEP1, REEP2, RTN2, SLC16A2, SLC33A1, SPART, SPAST, SPG7, SPG11, SPG21, TECPR2, TFG, TPP1, UBAP1, UCHL1, USP8, VPS37A, WASHC5, WDR48, ZFR, ZFYVE26 and ZFYVE27 |
| **Test seeds**<br>**(n=17)** | ACO2 (Bouwkamp *et al.*, 2018), ALS2 (Simone *et al.*, 2018), BICD2 (Kropatsch *et al.*, 2019), CCDC50, CCT5 (Bouhouche *et al.*, 2006), EXOSC3 (Blackstone, 2018a), GAD1 (Lo Giudice *et al.*, 2014), HACE1 (Akawi *et al.*, 2015), IFIH1 (Liu *et al.*, 2019), KCNA2 (Helbig *et al.*, 2016), KIDINS220 (Zhao *et al.*, 2019), LYST (Shimazaki *et al.*, 2014), MT-ATP6 (Verny *et al.*, 2011), MT-CO3 (Blackstone, 2018b), MT-ND4 (Pestronk, 2020), RETREG1 (Ilgaz Aydinlar *et al.*, 2014) and SELENOI (Ahmed *et al.*, 2017) |

Each protein of the global HSP-PPIN was scored based on the number of seeds to which it was directly connected, and a degree distribution was plotted (Supplementary Fig 2). All nodes interacting with at least 2 seeds (IIHs) were selected and used to extract the core HSP-PPIN composed of 164 nodes (including 45/57 HSP seeds [72.7%] and 8/11 test seeds [78.9%]) and 287 edges (Fig 1 and Supplementary File 2). The core HSP-PPIN represents the most interconnected part of the global HSP-PPIN graph and contains the interactors that are communal to 2 or more seeds, thus it can be used to investigate common functionalities across the different HSP genes.

**Figure 1.**
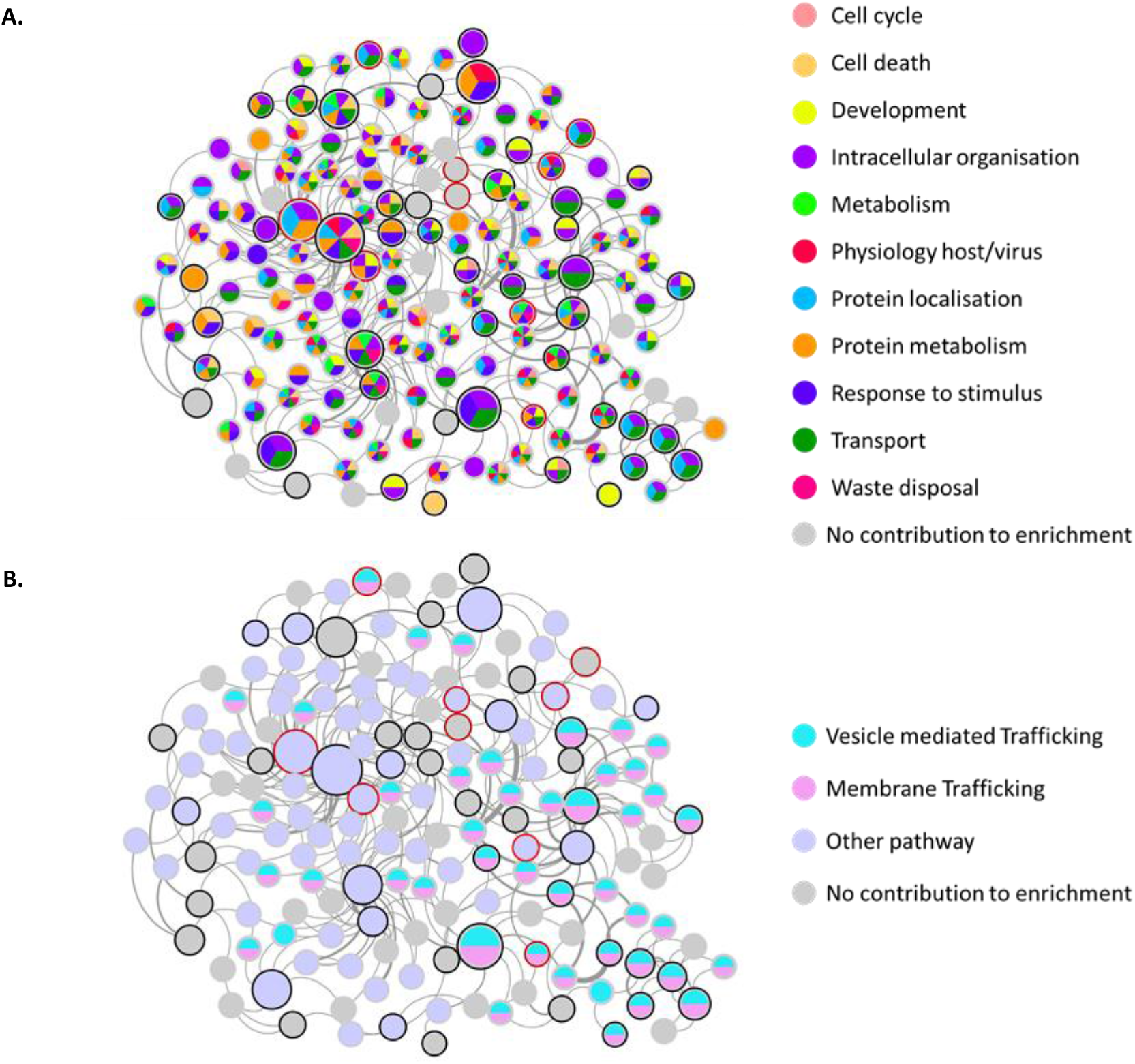
Functional enrichment of the core HSP-PPIN. The core HSP-PPIN is the most interconnected part of the global HSP-PPIN and includes i) the interactors connecting at least 2 seeds, and ii) the connected seeds. Seeds (HSP genes) are represented with a black border, test seeds with a red border (ACO2, ALS2, BICD2, CCDC50, CCT5, IFIH1, KIDINS220, LYST). The size of each node positively correlates with its number of connections (i.e. node degree) within the core HSP-PPIN. The thickness of each edge positively correlates with the final score of the respective interaction as calculated by PINOT (which is a proxy for confidence as it represents the sum of the number of different publications and number of different methods reporting the interaction). (**A**) Nodes contributing to the enrichment of functional blocks (built on Gene Ontology Biological Processes) are colour coded according to the legend (grey nodes are those that did not contribute to any of the enriched functional blocks). (**B**) The involvement of nodes of the core HSP-PPIN in pathways is visualised by node colour-coding based on Reactome’s pathway analysis.

Of note, the test-seed CCDC50 is present in the core HSP-PPIN and directly interacts with 2 proteins that are interactors of 6 HSP seeds. Comparatively, 95.5% of the proteins within the global HSP-PPIN, and 74.5% of the proteins within the core HSP-PPIN interacted with less than 6 HSP seeds. The strong connectivity of CCDC50 with HSP seeds indicates that they might be functionally related, and thus further supports the hypothesis that CCDC50 could be an HSP gene based on its genetic location [CCDC50 is located at 3q28 (https://www.ncbi.nlm.nih.gov/gene/152137), while the genetic loci of SPG14 is 3q27-28 (Boutry *et al*., 2019)].

### Functional enrichment: intracellular organization and trafficking

The nodes composing the core HSP-PPIN were analysed through functional enrichment to identify associated Gene Ontology Biological Processes (GO-BPs). Three different enrichment tools were used (g:Profiler, PantherGO and WebGestalt; Supplementary File 3). Despite p-values being corrected differently in the different tools, the enrichment ratio was calculated *via* the same formula (see Materials and Methods). We therefore selected the top 10 GO-BP terms (based on the enrichment ratio) from each of the 3 tools (Fig 2). The majority of the top-terms indicated functions such as those of “Transport” or “Intracellular organisation” (collectively accounting for 60-70% of terms significantly enriched using the 3 tools). The remaining terms referred to “Cell death” and “Physiology-host/virus” with important reference to protein targeting and the endomembrane system. Of note, we observed a 60% match of the GO-BPs in the top 10 enriched terms across all the 3 tools, and 60-100% match between at least 2 tools (g:Profiler: 100%, WebGestalt: 100%, and PantherGO: 60%). The unique terms from each tool, however, were closely related to already shared terms (e.g. “Anterograde axonal transport” [unique to PantherGO] is closely related to “Retrograde neuronal dense core vesicle transport” and “Retrograde axonal transport” [g:Profiler, WebGestalt and PantherGO]) (Fig 2).

**Figure 2.**
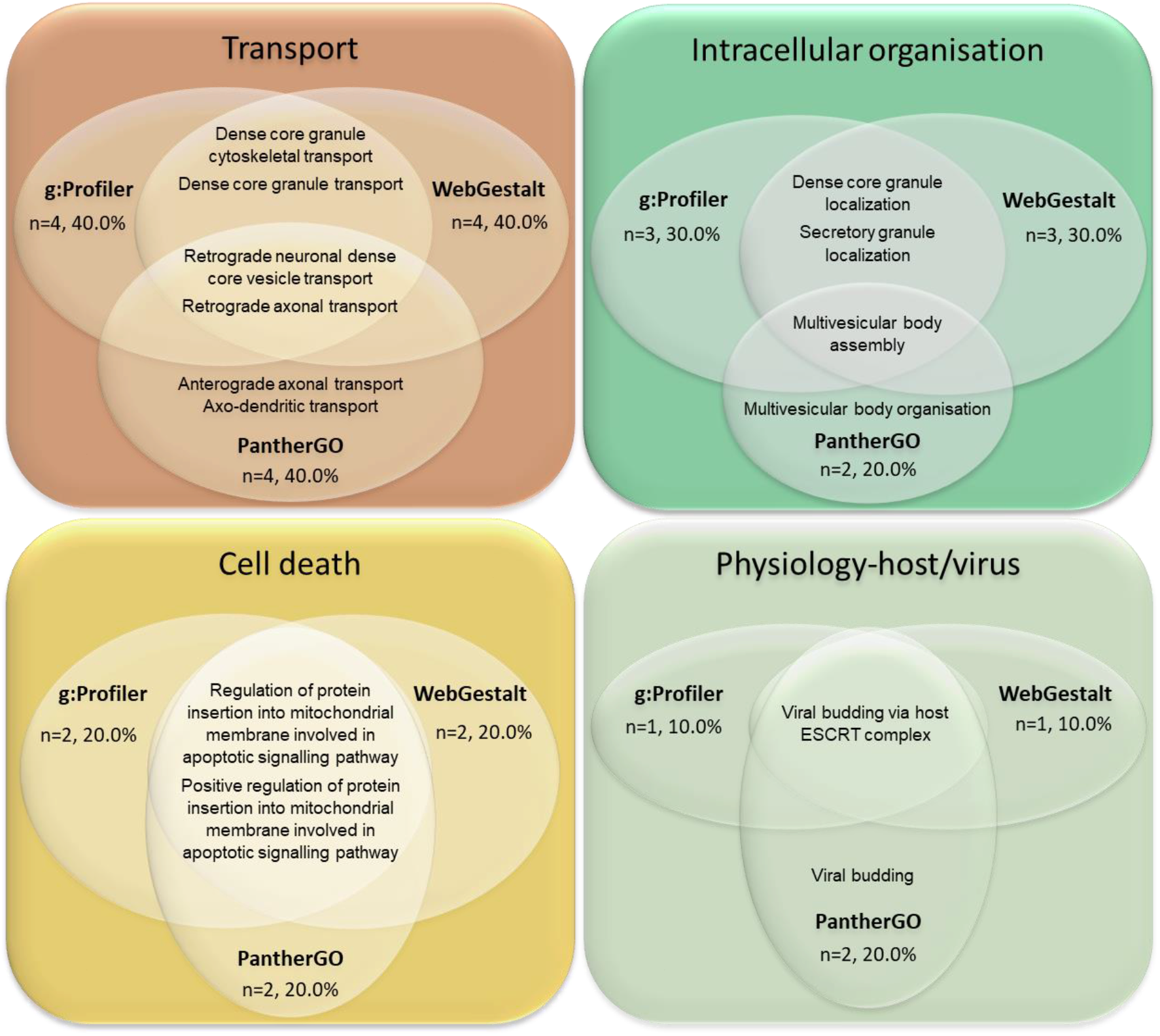
Top 10 GO-BPs enriched within the core HSP-PPIN. The 10 GO-BP terms from the functional enrichment of the core HSP-PPIN with the highest enrichment ratio were grouped into functional blocks based on semantic similarity. Most of the terms resulted from at least 2 enrichment tools (g:Profiler & WebGestalt: n=10/10, 100%; PantherGO: n=6/10, 60%).

The entirety of enriched GO-BP terms were then grouped by semantic similarity into semantic classes, which were further organised into functional blocks, thus aiding the interpretation of the enrichment results (see Materials and Methods and (Bonham *et al*, 2018; Ferrari *et al*, 2018; Ferrari *et al*, 2017; Tomkins *et al*, 2018)).

The raw results from each tool were similar in all 3 levels explored: the identity of the GO-BP terms, of the semantic classes and of the functional blocks. In fact, most of the GO-BP terms were common to at least 2 tools (n=115/171, 67.3%) (Supplementary Fig 3A), while after semantic classification of GO-BPs a higher proportion of semantic classes derived from at least 2 tools (n=49/58, 84.5%) (Supplementary Fig 3B). Finally, all the functional blocks were represented by all 3 tools (n=11/11, 100.0%) (Supplementary Fig 3C). Overall, this confirmed the consistency of results across different enrichment tools. However, these results also showed that even if consistency is very high at the more general levels of semantic classes and functional blocks, discrepancies can occur at the very specific GO-BP term level. Therefore, we decided to improve functional interpretation and reduce tool specific bias in further analyses by merging the GO-BP terms derived from the 3 tools within functional blocks replicated in at least 2 tools (in this case all terms) and adjusting the threshold of the p-value (see Materials and Methods).

The majority of significant GO-BP terms from the core HSP-PPIN enrichment analysis were associated with the functional block “Intracellular organisation” (22.2%), followed by “Transport” (19.3%), and then “Protein localisation” (13.5%), collectively accounting for more than half of GO terms (55.0%) (Fig 3, Supplementary Fig 4, Supplementary File 3). This result confirmed the findings previously obtained from the top-10 enriched terms, suggesting a role for these processes in the molecular mechanism(s) underlying HSP pathogenesis. Finally, and to increase specificity of the enrichment approach, we performed text mining for single words within all the significantly enriched GO-BP terms and detected significant enrichment for “axon” (n=7/171, 4.1% [8.9 fold enrichment] p<10^−10^ after 1000 random simulation), “endosomes” (n=3/171, 1.8% [5.7 fold enrichment] p<10^−10^), “membrane” (n=24/171, 14.0% [5.7 fold enrichment], p<10^−10^), “neurons” (n=9/171, 5.3% [3.4 fold enrichment], p=7.85 10^−7^), “projection” (n=6/171, 3.5% [5.4 fold enrichment], p=6.54 10^−7^), and “vesicles” (n=10/171, 5.8% [4.5 fold enrichment], p<10^−10^).

**Figure 3.**
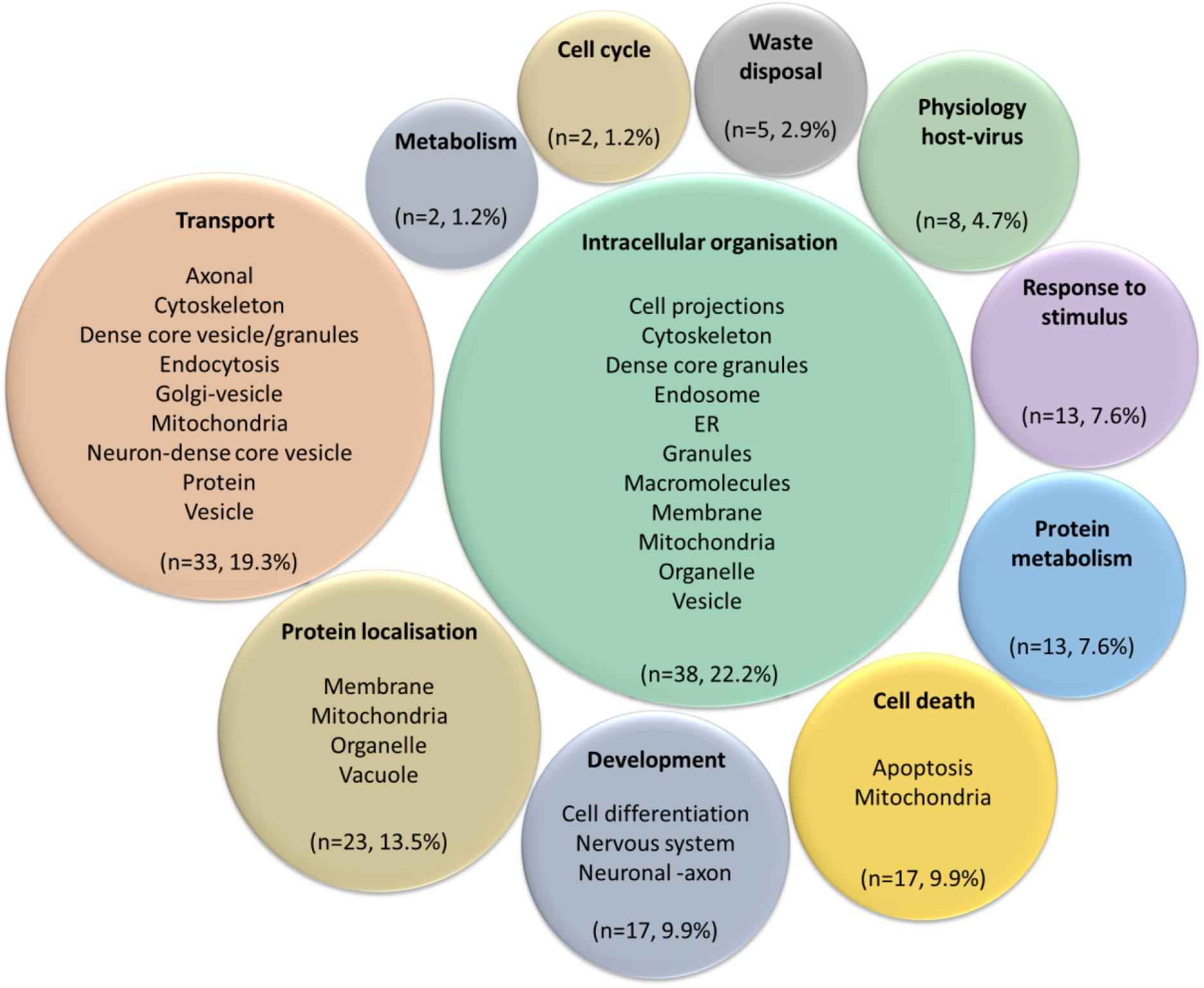
Graphical representation of the functional enrichment of the core HSP-PPIN. Functional enrichment was performed on the nodes of the core HSP-PPIN. The resulting GO-BP terms (n=171) (Supplementary File 4) were grouped into semantic classes (brief descriptions of several semantic classes are inside each circle) and then into functional blocks (title of each circle, bolded). The number and percentage of terms in each functional block was calculated for g:Profiler, WebGestalt, and PantherGO as described in Materials and Methods. For a more detailed version see Supplementary Fig 4.

Of note, the independent analysis of the core HSP-PPIN through Reactome (Supplementary File 3), suggested similar enrichment, whereby the 2 most significantly enriched pathways were: vesicle-mediated transport (REA identifier: R-HSA-5653656, p<10^−10^, 46 (28.0%) contributing nodes) and membrane trafficking (REA identifier: R-HSA-199991, p<10^−10^, 44 (29.3%) contributing nodes) (Fig 1B).

### Stratification of HSP clinical groups into 2 clusters

HSPs can present with a wide set of clinical features, with marked phenotypic heterogeneity between different patients. The complex forms of HSPs are defined by the co-occurrence of additional clinical features, the most frequently reported being: peripheral neuropathy (P), thinning of the corpus callosum (T), seizures (S), dementia or mental retardation (D) and optic atrophy (O). Finally, some patients also present with an early disease onset (E). Interestingly, medical reports and case studies sometimes state the presence of the above features in association with specific mutations in HSP genes. We have taken advantage of that this knowledge and grouped the genes based on the features with which they are associated. Therefore, the seeds within the core HSP-PPIN were coded based on their associated clinical features (Supplementary Fig 5). Of note, some seeds are associated with a single feature (n=9/57, 16%) while others are responsible for 2 (n=18/57, 32%), 3 (n=12/57, 21%) or 4 (n=7/57, 12%) clinical features. This seed characterisation allowed the extraction of 6 smaller subnetworks from the core HSP-PPIN, each of them containing the interconnected seeds (and their interactors) associated with each specific feature mentioned above (Supplementary Fig 6).

Enrichment of biological processes was performed on each clinical subnetwork separately, as previously described, using g:Profiler, PantherGO and WebGestalt (Supplementary File 4 and Supplementary Fig 7). The enrichment results obtained from the 3 tools were compared to assess their reproducibility and identify GO-BP terms of functional blocks that were replicated in at least 2 tools. These terms were merged to increase functional coverage as described above. The percentage of GO-BP terms within each functional block was calculated to weight its relevance. Principal components analysis (PCA) was then applied to reduce the complexity of the results obtained from the functional enrichment analyses to 2 principal components (PC1 and PC2). PCA thus allowed comparison of the 6 clinical subnetworks (Fig 4A). Interestingly, some of the clinical subnetworks functionally clustered together. Of note, this result was obtained with PCA performed on both the percentage of the GO terms in each functional block (Fig 4A) and their absolute numbers (Supplementary Fig 8A).

**Figure 4.**
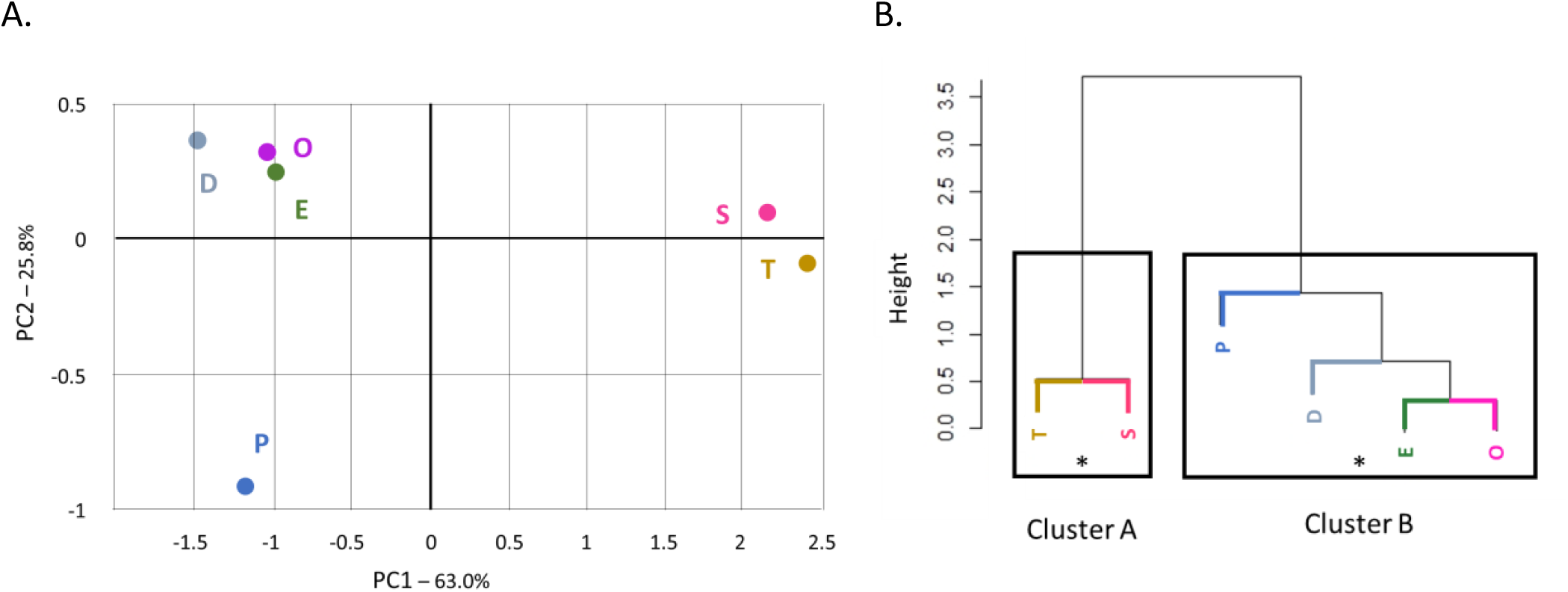
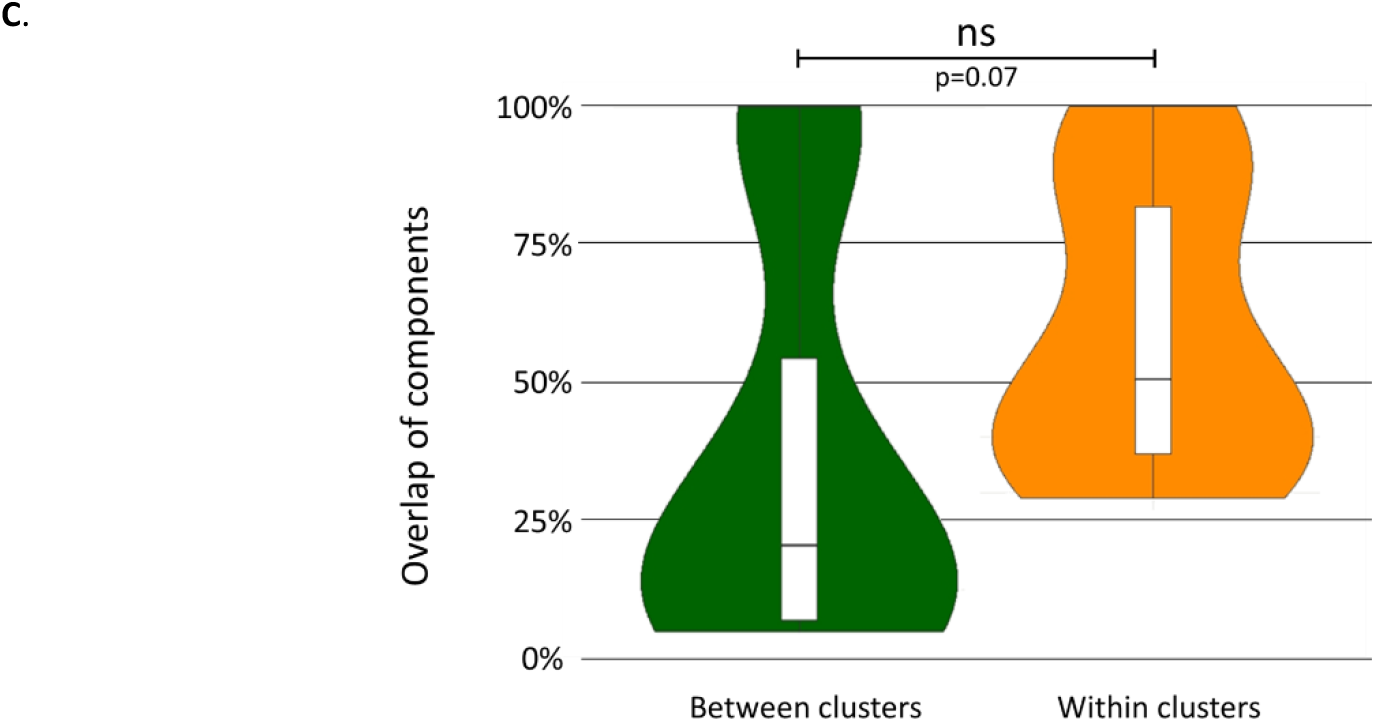
Comparison of the functional profiles of the 6 clinical subnetworks. (**A**) In the PCA graph each clinical subnetwork is represented by a single point of coordinates calculated based on PCA performed for the percentage of GO-BP terms and adjusted based on the explained variation of each axis (for details see Materials and Methods) [i.e. (x, y) = (PC1×0.630, PC2×0.258)]. (**B**) Cluster dendrogram produced based on hierarchical clustering of the gene groups as analysed in (A), in which the 2 suggested clusters are shown. *: pvclust-p-value>0.90 (pvclust-p-value A=0.99, pvclust-p-value B=0.91) E: Early onset, P: Peripheral neuropathy, T: Thin corpus callosum, S: Seizures, D: Dementia or mental retardation, O: Optic atrophy. (**C**) The percentage of protein identity between gene groups within the same cluster (EPOD and TS cluster) was compared to the protein identity between gene groups of different clusters using t-test (two-tailed, unequal distribution).

The PCA plot provided a first visual insight into potential functional clustering that was further confirmed by hierarchical clustering. Results were plotted into a cluster dendrogram (Fig 4B & Supplementary Fig 9) and the exact number of clusters to best fit the data was determined by 2 methods: Silhouette method and Multiscale bootstrap resampling (Supplementary Fig 10). Both methods suggested the presence of 2 clusters (named clusters A and B) in the cluster dendrogram (Silhouette method: the highest score was for 2 clusters; Multiscale bootstrap resampling: Cluster A and B had a pvclust-p-value=0.99 and 0.91, respectively, showing 99% and 91% confidence in the result). Cluster A is composed of thin corpus callosum, and seizures (thereafter named TS), while cluster B is composed of early onset, peripheral neuropathy, optic atrophy and dementia or mental retardation (thereafter named EPOD).

The co-clustering of the T and S subnetworks within the TS cluster is not surprising as they had 23 common proteins (n=23; T∩S = 82.1%, S∩T = 100%). However, we also observed a large overlap of proteins between the subnetworks of O and P (n=39; O∩P = 92.9%, P∩O = 53.4%); T and D (n= 25; T∩D = 89.3%, D∩T = 43.9%); S and E (n=23; S∩E = 100%, E∩S = 20.2%); and D and E (n=55; D∩E = 96.5%, E∩D = 48.2%). In all these cases, the common composition was large, yet not able to guide the order of similarity based on the dendrogram, nor to promote the co-clustering (Fig 4B). A full report of the overlaps between the clinical subnetworks is detailed in Supplementary Tables 3-5.

Plotting the percentages of overlaps across different clinical subnetworks allowed for running a statistical comparison. When considering the overlap of the subnetworks within cluster TS and within cluster EPOD (networks within the same cluster) in comparison with the overlaps of the subnetworks in TS vs EPOD (networks in different clusters) we found a non-significant difference in their distributions (p= 0.07; Fig 4C). This result suggests that the generation of the 2 distinct clinical clusters was highly affected by similarities in the functional profile of the subnetworks in terms of GO-BPs, while the overlap of nodes had a small or potentially no contribution.

### Differences between the clinical clusters based on functions and subcellular localisation

The potential differences of the 2 clinical clusters were further explored by performing enrichment analysis for GO-BPs using as input the protein components of the 2 clusters, TS and EPOD (Supplementary File 5). The comparison of the 2 obtained functional profiles is shown in Fig 5A and Supplementary Fig 11. Despite an overlap in the identity of the GO-BPs functional blocks between the 2 clusters (TS: n=4/5, 80%; EPOD: n=4/10, 40%), the granular distribution of specific GO-BP terms in each functional block differs between clusters, with the GO-BP functional blocks of: “Waste disposal” (+12.7-fold [compared to the core HSP-PPIN]), “Metabolism” (+9.3-fold), and “Protein metabolism” (+2.15-fold) being more represented in the TS rather than in the EPOD cluster (−0.13, −1.0, and 0.25-fold respectively) (Fig 5A). Meanwhile, the GO-BP functional blocks “Physiology-host/virus” (+0.22-fold), “Cell cycle” (+0.1-fold), and “Cell death” (+0.1-fold) were more represented in the EPOD rather than in the TS cluster (−1.0, −1.0, and −1.0-fold, respectively). Interestingly, 5 GO-BP terms related to the unfolded protein response (e.g. “Cellular response to unfolded protein” and “Cellular response to topologically incorrect protein”) were unique to the TS cluster (n=5/25, 25%), even with cluster EPOD having a 6-fold higher number of total GO-BP terms (n_GO-BPtotalEPOD_=158 vs N_GO-BPtotalTS_=25), thus highlighting the importance of protein folding for the TS cluster only. Overall, these results of GO-BP enrichment indicated that functions associated with protein metabolism, waste disposal and unfolded protein response might be more important processes in the TS rather than in the EPOD cluster; while the EPOD cluster presents with a functional enrichment profile very similar to that of the entire core HSP-PPIN.

**Figure 5.**
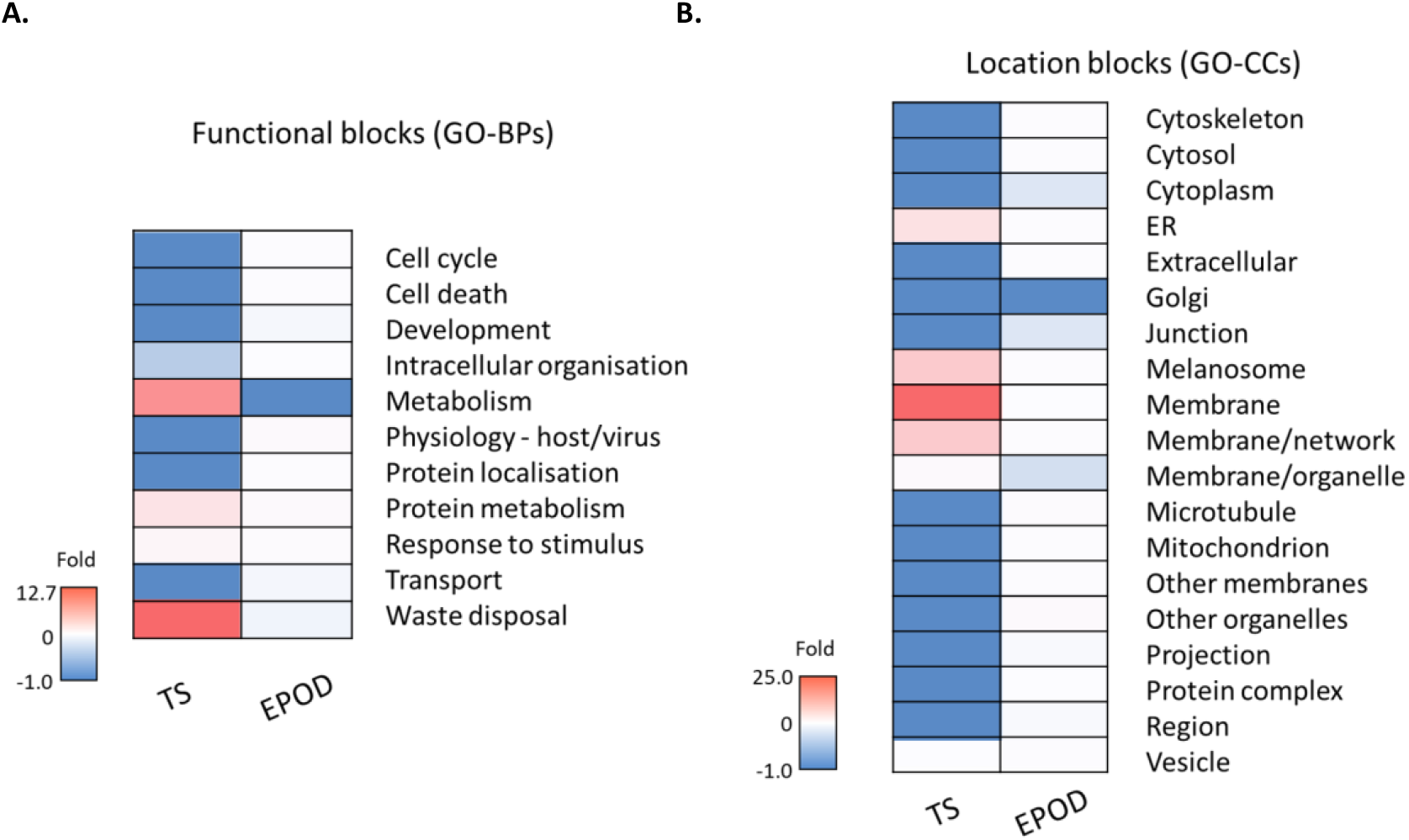
Differential patterns of enrichment for the TS and EPOD clusters. The distribution of the GO-BPs (**A**) and GO-CCs (**B**) of the clusters, TS and EPOD, are presented as a fold change compared to the profile of the core HSP-PPIN. A more detailed version is shown in Supplementary Fig 11, while the totality of the results is shown in Supplementary File 5.

Similarly, we performed Gene Ontology Cellular Component (GO-CC) enrichment using as input the protein components of the 2 clusters A and B (Supplementary File 5). This analysis showed additional differences between the profiles of the 2 clusters (Fig 5B, Supplementary Fig 11). Even though, there are common GO-CC location blocks between the 2 clusters (TS: n=5/6, 83.3%; EPOD:, n=5/17, 29.4%), their order of relevance based on the percentage of GO-CC terms differed substantially. Interestingly, and confirming the results obtained previously with GO-BPs, a higher percentage of GO-CC location blocks are related to “ER” (+4.7-fold [compared to the core HSP-PPIN]), “Melanosomes” (+8.5-fold), and “Membranes” (i.e. “Membranes”: +25.0-fold, “Membrane/network” +8.5-fold, and “Membranes/organelle” +0.5-fold) for the TS cluster in comparison with the EPOD cluster (0.13, 0.13, 0, 0.13, −0.30-fold, respectively). As for the EPOD clusters, higher relevance is observed in the GO-CC location blocks: “Other organelles” (+0.5-fold), “Microtubules” (+0.4-fold), “Cytoskeleton”, “Cytosol”, “Extracellular”, ““Mitochondria” and “Other membranes” (+0.1-fold) than in TS (−1-fold in TS).

## Discussion

Network-based approaches have been increasingly used to study complex human diseases, such as neurodegenerative diseases and cancer (Manzoni *et al*., 2020). The Hereditary Spastic Paraplegias (HSPs) are neurodegenerative diseases with considerable genetic and clinical heterogeneity (Boutry *et al*., 2019; Faber *et al*., 2017), rendering them particularly interesting to study using a protein-protein interaction network (PPIN) approach. We applied a bottom-up approach, starting with the selection of genes involved in the disease and built the relevant interactome around them. We focused on experimentally validated human PPIs of HSP genes, not including genes associated with a disease spectrum in which HSP is involved (e.g. HSP-ataxia spectrum) or genes with related phenotype, in contrast with prior studies (Bis-Brewer *et al*, 2019; Novarino *et al*, 2014; Parodi *et al*, 2018; Synofzik & Schule, 2017). While the excluded data might be useful in the effort to conceptualise the possible interactions and mechanisms of HSP related diseases, they were not considered to be specific or supported strongly enough to be included in our analysis.

We applied the PINOT pipeline to mine the curated literature and download PPIs for each single seed, thus obtaining each seed’s interactome (Ferrari *et al*., 2018). We then constructed the global HSP-PPIN by combining each seed’s interactome in a modular fashion. We finally filtered the global HSP-PPIN, excluding the nodes that interacted with a single seed, thus retaining those interactors that were bridging at least 2 seeds’ interactomes. This step allowed for removal of all the unique interactors of each seed and for the extraction of the core HSP-PPIN, which is the most connected part of the network, containing nodes that are shared across seeds, and responsible for connections across different interactomes. By containing all the shared interactors and connections among seeds, the core HSP-PPIN can be used to infer shared functions communal to multiple HSP genes. (Tomkins *et al*., 2020).

It is important to observe that most HSP seeds are indeed part of the core HSP-PPIN, meaning they are connected through at least one shared interactor. This result suggests that they are likely to be functionally related (based on the guilt-by-association principal (Oliver, 2000)) and therefore convergent molecular mechanism(s) drive disease pathogenesis, regardless of the mutated gene acting to initiate the degenerative process. The seeds that were absent from the core HSP-PPIN (i.e. seeds that do not share any interactors with other seeds) had a low number of curated interactors ranging from 0 to 4 (*PLA2G6, CPT1C, CYP2U1, C12orf65, B4GALNT1, TECPR2, ENTPD1, ATL1, SPG11, DDHD1, AP5Z1, SLC16A2, GAD1, RAB3GAP2, and HACE1*). With limited interactors, their absence from the core HSP-PPIN could be the result of ascertainment bias (i.e. these seeds are understudied proteins with limited number of known interactors) rather than representing a more fundamental divergence in aetiology (Schaefer *et al*, 2015) As more PPIs are discovered, the human interactome will become more complete (Huttlin *et al*, 2017; Luck *et al*, 2020; Rolland *et al*, 2014; Wewer Albrechtsen *et al*, 2018) and might be able to help us better understand the connecting processes of large groups of genes and potentially point towards the disease mechanism. Exceptions were EXOSC3 (test-seed), SPG21 (HSP-seed) and KCNA2 (test-seed) with 21, 10 and 6 interactors, respectively. In this second scenario, it can be hypothesised that these seeds are not interacting with other HSP seeds, meaning that, by not sharing the same interactome, they might potentially be associated with different molecular mechanisms of disease.

In this study we included 17 test seeds, genes for which there is no clear consensus regarding their potential association with HSPs, as they have been controversially reported in clinical literature. Eight of the test seeds (i.e. ALS2, BICD2, CCDC50, CCT5, KIDINS220, ACO2, LYST and IFIH1) were present in the core HSP-PPIN, providing *in silico* evidence of their relevance within the HSP protein interaction landscape. The presence of five of those test seeds (i.e. CCT5, KIDINS220, ACO2, LYST and IFIH1) correlates with the processes and cellular components indicated to play a role in HSPs from previous and the current work, namely of lysosomal homeostasis, protein folding and transport, cell death, neurodegeneration, and antiviral responses, with which they also have been associated (Crow *et al*, 2020; Faigle *et al*, 1998; Freund *et al*, 2014; Leong & Chow, 2006; Liao *et al*, 2007; Spiegel *et al*, 2012). The presence of ALS2 in the core HSP-PPIN is not surprising, as it is considered an HSP gene by many clinicians and researchers (Boutry *et al*., 2019; de Souza *et al*, 2017; Lo Giudice *et al*, 2014). An interesting test-seed present in the core HSP-PPIN is CCDC50, because it was included in this study based on its chromosomal location being within the locus of SPG14 [CCDC50 is located at 3q28 (https://www.ncbi.nlm.nih.gov/gene/152137), while the genetic loci of SPG14 is 3q27-28 (Boutry *et al*., 2019)).). Of note, CCDC50 formed interactions with more seeds than most interactors of the global HSP-PPIN and the core HSP-PPIN. This result represents an *in silico* prediction that alterations in CCDC50 could be leading to the HSP type SPG14 and it suggests to include CCDC50 in the list of prioritized genes to be screened for rare variant discovery.

Notably, the protein product of the gene *RNF170* was found to be associated with HSPs (and published) after this analysis commenced (Wagner *et al*., 2019) and was indeed present within the global HSP-PPIN. This result demonstrates the utility in using PPINs to study complex disorders, as they can aid prioritisation of candidate genes from genetic analysis (Erlich *et al*, 2011) and hint to key proteins involved in disease mechanisms.

The analysis of a disease-focused PPIN based on functional annotation provides an opportunity to gain a deeper understanding of the underlying mechanism(s) of disease using a holistic view (Koh *et al*., 2012). Therefore, enrichment analysis was performed for the components of the core HSP-PPIN, supporting the involvement of some of the processes previously suggested to be associated with the disease mechanism of HSPs. Out of the 10 mechanisms suggested by Lo Giudice et al (Lo Giudice *et al*., 2014), 3 were supported by the results of this work were 3, namely, “endosome membrane trafficking and vesicle formation”, “abnormal membrane trafficking and organelle shaping”, “dysfunction of axonal transport”, but also, 3 additional processes, namely, “autophagy”, “axon development” and “abnormal cellular signalling in protein morphogenesis”, while we did not find evidence in our analysis for “oxidative stress”, “abnormal lipid metabolism”, “abnormal DNA repair” and “dysregulation of myelination”. Regarding the mechanisms hypothesised by de Souza and colleagues (de Souza *et al*., 2017), those in accordance with this work were “intracellular active transport”, “endolysosomal trafficking pathways” and “ER shaping”, while we did not find evidence in our analysis for “lipid metabolism”, “mitochondrial dysfunction”, nor “migration and differentiation of neurons”. Our results are more in line with the suggestion from Blackstone (Blackstone, 2018a) that the key biological processes at play in the aetiopathogenesis of HSPs are “organelle shaping and biogenesis” and “membrane cargo and trafficking”, further supporting the notion that HSPs could be considered transportopathies (Gabrych *et al*, 2019), and that the dysregulation of ER morphology and function could be implicated in HSPs (Lee & Blackstone, 2020). However, some of the suggested hypotheses, namely “nucleotide metabolism”, “mitochondrial function” and “lipid/cholesterol metabolism” (Blackstone, 2018a), were not supported by the findings of this study. Interestingly, functional data were not used for the creation of the HSP-PPINs, therefore the conclusions obtained here are only based on PPIs and represent a further validation of some of the published functional analyses. These results highlight the potential of a PPIN analysis approach combined with functional enrichment to identify the most relevant functions among the genes of interest related to a complicated disease, which is an important step for discovering disease modifying agents.

In order to explore if the clinical diversity of the HSPs reflects a mechanistic heterogeneity of disease, machine learning tools (PCA and hierarchical clustering) were used to analyse the functional profile of the core HSP-PPIN. Based on our *in silico* analysis, we suggest the existence of at least 2 main subtypes of HSPs. The first functional subtype includes the clinical features of thin corpus callosum and seizures (i.e. TS cluster); while the second gathers those cases characterized by early onset, peripheral neuropathy, dementia or mental retardation and optic atrophy (i.e. EPOD cluster). Further analysis for biological processes of the 2 clinical clusters suggested that “protein metabolism” and “waste disposal” are prominent in the TS cluster. In addition, most of the unique results for this cluster were related to the unfolded protein response. These results support the relevance of the regulation of protein level and conformation for the TS cluster. While for the EPOD cluster, the most important functions were related to “physiology-host/virus” and “cell death”, which suggest that the endomembrane system involved in the viral process, together with mechanisms involved in cell survival are of higher importance in the EPOD cluster.

These findings were further supported by cellular component and pathways analysis, where the TS cluster showed a higher enrichment in different types of membranes, melanosomes and the ER, while results for the EPOD cluster were more focused on extracellular components, mitochondria, other organelles and the cytoskeleton.

The results presented in this study require further functional validation, however they provide a platform indicating that HSP patients could be stratified based on the molecular mechanisms involved in disease aetiopathogenesis and this in turn can be beneficial for developing therapeutic strategies and aiding efforts to stratify patients for clinical trials.

This application provides insight into the utility of PPIN analysis in the study of complex disorders, as PPINs are a powerful tool that can extract and combine a large extent of previous data in a relatively quick and easy fashion. Using this approach can create a comprehensive picture that summarises the current knowledge, helping in prioritising and confirming existing mechanistic theories, guiding research based on the identification of interesting proteins and pathways, as well as highlighting uncertain areas that require further investigation.

## Materials and Methods

### Selection of seeds

The protein products of 83 genes were selected as seeds based on their clinical relevance for HSPs (de Souza *et al*., 2017), among which 16 have not been widely recognised as HSP genes hereafter referred to as test seeds (Table 1 and Supplementary Table 2).

### Collection of PPIs and HSP-PPINs

The 83 seeds were used as the input to query the PINOT webtool (Tomkins *et al*., 2020) [http://www.reading.ac.uk/bioinf/PINOT/PINOT_form.html]. PINOT produces a list of experimentally demonstrated binary PPIs containing unique, human PPI data obtained by merging and processing PPI data from 7 databases: BioGrid (Oughtred *et al*, 2019), InnateDB (Breuer *et al*, 2013), IntAct (Orchard *et al*, 2014), MBInfo (Singapore, 2019), MINT (Licata *et al*, 2012), UniProt (UniProt, 2019) and bhf-ucl. Through PINOT, interactions are filtered and scored based on the number of publications that report a particular interaction and the number of different methods used for their detection. The interactions provided from PINOT were then screened to remove PPIs with a final score <3 (those interactions without replication in the curated literature). The retained interactions were visualised using Cytoscape (v3.7.2), thus creating the global HSP-PPIN.

Each node in the network was scored based on the number of seeds to which it connected. The nodes interacting with more than one seed, referred to as “inter-interactomes hubs (IIHs)” (Ferrari *et al*., 2017), were used to extract a subnetwork composed of IIHs and the connected seeds. This subnetwork was termed the “core” HSP-PPIN.

The interactions for the global HSP and core HSP networks were downloaded on the 09/07/2019, PINOT (beta version), using the stringent and *Homo sapiens* filters (default).

### Enrichment analyses

The subset of proteins composing the core HSP network underwent enrichment analysis (Biological Processes [BPs] and/or Cellular Components [CCs] Gene Ontology [GO] annotations). The consistency of the results was evaluated by using 3 independent online tools, which utilise different algorithms, multiple test correction and/or versions of the GO database. In particular: g:Profiler (July 2019, Over-representation enrichment analysis (Fisher’s one tailed test), Bonferroni’s corrections, GO database release 11/07/2019, excluding electronic annotations and analysed against the annotated human genome) (Reimand *et al*, 2016) [https://biit.cs.ut.ee/gprofiler/gost], Gene Ontology using Panther’s tool (September/October 2019, Binomial test, Bonferroni’s corrections, GO database release 03/07/2019, analysed against the human genome) (Ashburner *et al*, 2000; Mi *et al*, 2017; The Gene Ontology, 2019) [http://geneontology.org/ and http://pantherdb.org/] and WebGestalt (October 2019, Over-representation enrichment analysis (Hypergeometric test), FDR, GO database release 14/01/2019, analysed against the protein coding human genome) (Wang *et al*, 2017) [http://www.webgestalt.org/].

The output of the functional enrichment includes a list of enriched GO terms and their respective enrichment ratio which can be calculated using the following formulas:

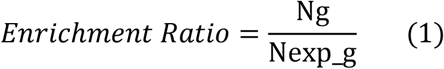

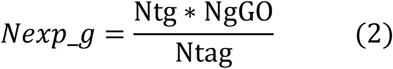

where Ng is the number of genes with a GO term in the data, Nexp_g the number of expected genes with a GO term in the data, Ntg the number of genes in the data, NgGO the number of genes annotated with a GO term in the GO database, and Ntag the total number of annotated genes in the GO database.

The enriched BP and CC GO terms were grouped by semantic similarity into semantic classes using in-house developed dictionaries. The semantic classes were further clustered into functional blocks and location blocks, respectively. The GO terms classified in the semantic classes “general” and “metabolism” were not included in the analysis as they refer to GO terms that provide limited functional specificity to the analysis (Ferrari *et al*., 2017).

Finally, in order to reduce any tool specific bias, only the functional or location blocks confirmed to be enriched by at least 2 of the 3 enrichment tools (g:Profiler, PantherGO and WebGestalt) were retained for further analysis. Particularly, for those blocks that were replicated across at least 2 tools, we analysed the merge of their semantic classes resulting from each individual tool. The threshold for determining statistical significance of each GO term was therefore decreased to p=0.0166 (=0.05/3). Additionally, only the terms that were enriched in association with at least 4 genes were retained.

The comparison of the clusters’ enrichment profiles for BP and CC was performed by calculating the following ratio for each block:

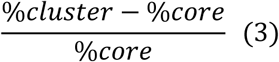

where %cluster is the percentage of GO terms of a cluster, and %core is the percentage of GO terms of the core-HSP-PPIN.

In the case that the aforementioned ratio of the functional or location block had the value of zero for the core dataset, since dividing by zero results to ∞, we set up 25 as the maximum value and −25 as the minimum value for visualisation purposes.

Pathway enrichment was performed using Reactome’s online analysis tool (v69 & v70 in September and December 2019) (Jassal *et al*, 2020) [https://reactome.org/PathwayBrowser/#TOOL=AT]. The pathways that were significantly enriched (p-value<0.05) were retained and filtered further to remove those with 3 or less proteins involved.

Text mining was performed on the GO-BP terms after the merging of results from the 3 tools. The number of terms related to axons, cytoskeleton, endosomes, membranes, neurons, projections and vesicles were counted based on the presence of “axo*”, “cytoskelet*”, “endos*”, “membrane*” “microtubu*”, “vesic*”, “neuro*” and “projections*”, respectively. An enrichment analysis was performed using the same key words, based on their frequency in the results versus in the in-house dictionary that included a collection of GO terms, using the described formulas (1) & (2). The statistical analysis was performed by running 100,000 random simulations where these key words were extracted from the in-house dictionary, and the pnorm() value was calculated.

### Principal component analysis & Hierarchical clustering

In order to compare functional enrichment profiles, Principal Component Analysis (PCA) was conducted through R (v. 4.0.2) using the prcomp() function of the stats package. The analysis of the number and percentage of GO terms in each functional block were both rendered necessary due to the substantial difference in the number of resulting GO terms of the 6 groups, whose functional enrichment profiles were compared (22<n<114) (Supplementary File 4).

Hierarchical clustering was performed using the hclust() function (R stats package) for the groups in the PCA plot, using Euclidean as a distance measure for row clustering. However, one unit of distance in the x axis of the PCA plot is more important than on the y axis, due to PC1 (x axis) explaining more variation than PC2 (y axis) (63% and 25.8%, respectively for the analysis based on the percentage of GO terms). Thus, the coordinates of each point had to be transformed; they were multiplied by the explained variation, so that the distance between points can have the same significance in any direction and can thus be used for hierarchical clustering. Through Hierarchical clustering, the cluster dendrogram was produced. Choosing the best fit for the number of clusters derived from Hierarchical clustering was based on the Silhouette method (P.J., 1987) and the Multiscale bootstrap resampling method (Suzuki & Shimodaira, 2006). For the former, the index/score were calculated for 2 up to 6 clusters. The latter was based on the R package “pvclust” that assigns pvclust p-values to each branch of the dendrogram, which show the confidence of the result (the higher the value, the more confident we are of the result) (Suzuki & Shimodaira, 2006) (Supplementary File 1).

## Funding

NV is supported by the Engineering and Physical Sciences Research Council grant to PAL [grant number EP/M508123/1].

JET is supported by the Biomarkers Across Neurodegenerative Diseases Grant Program 2019, BAND3 (Michael J. Fox Foundation, Alzheimer’s Association, Alzheimer’s Research UK and Weston Brain Institute [grant number 18063 awarded to CM and PAL]).

EK is the recipient of an HFSP long term fellowship (LT001044/2017).

PAL and JH are supported by the ASAP research network, grant number ASAP0478. PAL is supported by the Michael J. Fox Foundation. This work was further supported by the Medical Research Council [grant numbers MR/N026004/1 to JH and PAL; MR/L010933/1 to PAL]; the Wellcome Trust to JH [grant number 202903/Z/16/Z; the National Institute for Health Research University College London Hospitals Biomedical Research Centre to JH; the UK Dementia Research Institute (which receives its funding from DRI Ltd, funded by the UK Medical Research Council, Alzheimer’s Society and Alzheimer’s Research UK); by the Dolby Family Fund; and by the BRCNIHR Biomedical Research Centre at University College London Hospitals NHS Foundation Trust and University College London.

## Author contributions

NV, CM and PAL set up the experimental protocol; NV, run the analyses; EK and HH assisted with the clinical aspects of the manuscript; JET assisted with PPI analysis; JH, MJT, CM and PAL provided critical supervision; NV and CM wrote the manuscript which was reviewed by all authors.

## Competing Interest

The authors declare that they have no competing interests

## List of Supplementary Material

### Supplementary Figures and Tables

**Supplementary File 1**: Interactions used for the creation of the global HSP-PPIN

**Supplementary File 2**: Names of components of the core HSP-PPIN and whether they are an HSP seed or test seed

**Supplementary File 3**: Enrichment data of the core HSP-PPIN

**Supplementary File 4**: Enrichment data of the clinical groups within the core HSP-PPIN

**Supplementary File 5**: Enrichment data of the clinical clusters within the core HSP-PPIN

## Notes

### Competing Interest Statement

The authors have declared no competing interest.

