## Supplementary Tables and Figures for "Integrating protein networks and machine learning for disease stratification in the Hereditary Spastic Paraplegias"

| Supplementary Table 1. Sources of variation in the clinical phenotype of HSP patients | |
|---|---|
| **Age of onset** | Early childhood up to late adulthood |
| **Form** | Pure and complex |
| **Mode of inheritance** | Autosomal recessive, autosomal dominant, X-linked, mitochondrial and unknown |
| **Usual Symptoms** | Bilateral spasticity and weakness of the lower body, leg hypertonicity, positive Babinski sign, muscle weakness, hyperreflexia, bladder dysfunction, loss of vibration sensation in the ankles, and pes cavus |
| **Additional symptoms present in some complex forms** | Cerebellar ataxia, epilepsy, cognitive or mental impairment, cataracts, retinal alteration, optic atrophy, peripheral neuropathy, dystonia, and parkinsonism |

**Supplementary Table 2. The gene (or locus) and protein name responsible for each HSP type**

| Gene name (or locus) | Protein name (previous name) | UniProt identifier (SwissProt) | HSP type |
|---|---|---|---|
| L1CAM | neural cell adhesion molecule L1 | P32004 | SPG1 |
| PLP1 | myelin proteolipid protein | P60201 | SPG2 |
| ATL1 | atlastin-1 | Q8WXF7 | SPG3A |
| SPAST | spastin | Q9UBP0 | SPG4 |
| CYP7B1 | 25-hydroxycholesterol 7-alpha-hydroxylase | O75881 | SPG5A |
| NIPA1 | magnesium transporter NIPA1 | Q7RTP0 | SPG6 |
| SPG7 | paraplegin | Q9UQ90 | SPG7 |
| WASHC5 | WASH complex subunit 5 (strumpellin) | Q12768 | SPG8 |
| ALDH18A1 | delta-1-pyrroline-5-carboxylate synthase | P54886 | SPG9A/SPG9B |
| KIF5A | kinesin heavy chain isoform 5A | Q12840 | SPG10 |
| SPG11 | spatacsin | Q96JI7 | SPG11 |
| RTN2 | reticulon-2 | O75298 | SPG12 |
| HSPD1 | 60 kDa heat shock protein, mitochondrial | P10809 | SPG13 |
| (3q27-q28) | | | SPG14 |
| ZFYVE26 | zinc finger FYVE domain-containing protein 26 (spastizin) | Q68DK2 | SPG15 |
| (Xq11.2) | | | SPG16 |
| BSCL2 | seipin | Q96G97 | SPG17 |
| ERLIN2 | erlin-2 | O94905 | SPG18/SPG37 |
| (9q33-q34) | | | SPG19 |
| SPART | spartin | Q8N0X7 | SPG20 |
| SPG21 | maspardin | Q9NZD8 | SPG21 |
| SLC16A2 | monocarboxylate transporter 8 | P36021 | SPG22 |
| DSTYK | dual serine/threonine and tyrosine protein kinase | Q6XUX3 | SPG23 |
| (13q14) | | | SPG24 |
| (6q23-24.1) | | | SPG25 |
| B4GALNT1 | beta-1,4 N-acetylgalactosaminyltransferase 1 | Q00973 | SPG26 |

| Supplementary Table 2. (continued) The gene (or locus) and protein name responsible for each HSP type | | | |
|---|---|---|---|
| **Gene name (or locus)** | **Protein name (previous name)** | **UniProt identifier (SwissProt)** | **HSP type** |
| (10q22.1-q24.1) | | | SPG27 |
| DDHD1 | phospholipase DDHD1 | Q8NEL9 | SPG28 |
| (1p31.1-p21.1) | | | SPG29 |
| KIF1A | kinesin-like protein KIF1A | Q12756 | SPG30 |
| REEP1 | receptor expression-enhancing protein 1 | Q9H902 | SPG31 |
| (14q12-q21) | | | SPG32 |
| ZFYVE27 | protrudin | Q5T4F4 | SPG33 |
| (Xq24-q25) | | | SPG34 |
| FA2H | fatty acid 2-hydroxylase | Q7L5A8 | SPG35 |
| (12q23-q24) | | | SPG36 |
| (4p16-p15) | | | SPG38 |
| PNPLA6 | neuropathy target esterase | Q8IY17 | SPG39 |
| (11p14.1-p11.2) | | | SPG41 |
| *SLC33A1* | acetyl-coenzyme A transporter 1 | O00400 | SPG42 |
| *C19orf12* | protein C19orf12 | Q9NSK7 | SPG43 |
| *GJC2* | gap junction gamma-2 protein | Q5T442 | SPG44 |
| *NT5C2* | cytosolic purine 5'-nucleotidase | P49902 | SPG45/SPG65 |
| *GBA2* | non-lysosomal glucosylceramidase | Q9HCG7 | SPG46 |
| *AP4B1* | AP-4 complex subunit beta-1 | Q9Y6B7 | SPG47 |
| *AP5Z1* | AP-5 complex subunit zeta-1 | O43299 | SPG48 |
| *TECPR2* | tectonin beta-propeller repeat-containing protein 2 | O15040 | SPG49 |
| *AP4M1* | AP-4 complex subunit mu-1 | O00189 | SPG50 |
| *AP4E1* | AP-4 complex subunit epsilon-1 | Q9UPM8 | SPG51 |
| *AP4S1* | AP-4 complex subunit sigma-1 | Q9Y587 | SPG52 |
| *VPS37A* | vacuolar protein sorting-associated protein 37A | Q8NEZ2 | SPG53 |
| *DDHD2* | phospholipase DDHD2 | O94830 | SPG54 |
| *C12orf65* | probable peptide chain release factor C12orf65, mitochondrial | Q9H3J6 | SPG55 |
| *CYP2U1* | cytochrome P450 2U1 | Q7Z449 | SPG56 |
| *TFG* | protein TFG | Q92734 | SPG57 |

| **Supplementary Table 2. (continued) The gene (or locus) and protein name responsible for each HSP type** | | | |
|---|---|---|---|
| **Gene name (or locus)** | **Protein name (previous name)** | **UniProt identifier (SwissProt)** | **HSP type** |
| *KIF1C* | kinesin-like protein KIF1C | O43896 | SPG58 |
| *USP8* | ubiquitin carboxyl-terminal hydrolase 8 | P40818 | SPG59 |
| *WDR48* | WD repeat-containing protein 48 | Q8TAF3 | SPG60 |
| *ARL6IP1* | ADP-ribosylation factor-like protein 6-interacting protein 1 | Q15041 | SPG61 |
| *ERLIN1* | erlin-1 | O75477 | SPG62 |
| *AMPD2* | AMP deaminase 2 | Q01433 | SPG63 |
| *ENTPD1* | ectonucleoside triphosphate diphosphohydrolase 1 | P49961 | SPG64 |
| *ARSI* | arylsulfatase I | Q5FYB1 | SPG66 |
| *PGAP1* | GPI inositol-deacylase | Q75T13 | SPG67 |
| *KLC2* | kinesin light chain 2 | Q9H0B6 | SPG68 |
| *RAB3GAP2* | rab3 GTPase-activating protein non-catalytic subunit | Q9H2M9 | SPG69 |
| *MARS* | methionine--tRNA ligase, cytoplasmic | P56192 | SPG70 |
| *ZFR* | zinc finger RNA-binding protein | Q96KR1 | SPG71 |
| REEP2 | receptor expression-enhancing protein 2 | Q9BRK0 | SPG72 |
| *CPT1C* | carnitine O-palmitoyltransferase 1, brain isoform | Q8TCG5 | SPG73 |
| *IBA57* | putative transferase CAF17, mitochondrial | Q5T440 | SPG74 |
| *MAG* | myelin-associated glycoprotein | P20916 | SPG75 |
| *CAPN1* | calpain-1 catalytic subunit | P07384 | SPG76 |
| *FARS2* | phenylalanine--tRNA ligase, mitochondrial | O95363 | SPG77 |
| *ATP13A2* | cation-transporting ATPase 13A2 | Q9NQ11 | SPG78 |
| *UCHL1* | ubiquitin carboxyl-terminal hydrolase isozyme L1 | P09936 | SPG79 |
| *UBAP1* | ubiquitin-associated protein 1 | Q9NZ09 | SPG80 |
| *TPP1* | tripeptidyl-peptidase 1 | O14773 | - |

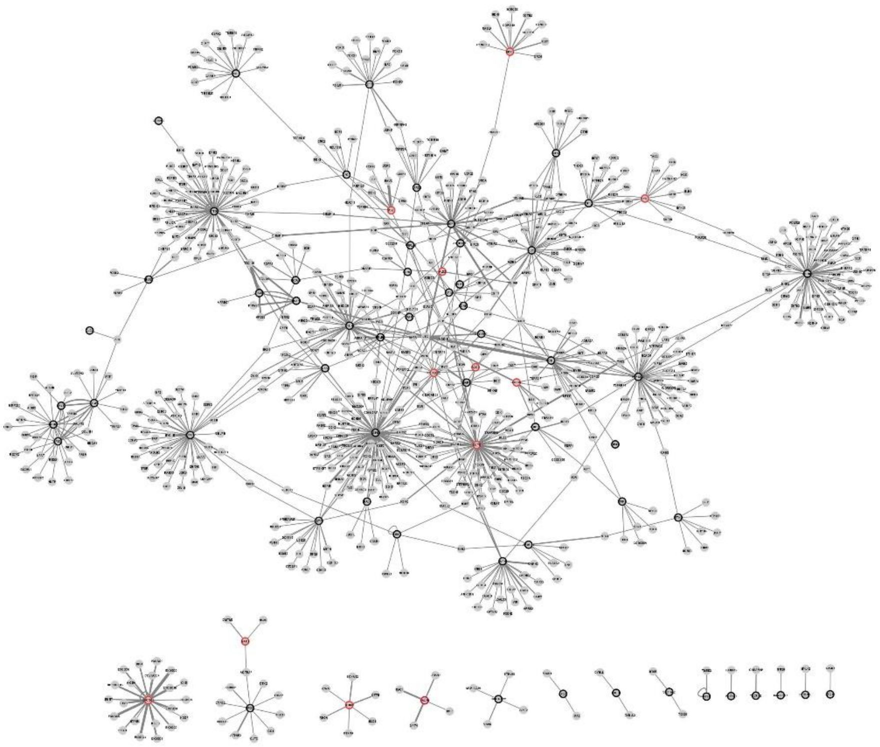

**Supplementary Figure 1. The global HSP-PPIN**
The global HSP-PPIN is a visualisation of all binary interactions of the HSP seeds and test seeds that were collected through the online tool PINOT following filtering based on the final score. The nodes corresponding to the HSP seeds have a black border, while the test seeds have a red border. The thickness of each edge positively correlates with the final score of the respective interaction as calculated by PINOT, which acts as a proxy for interaction confidence.

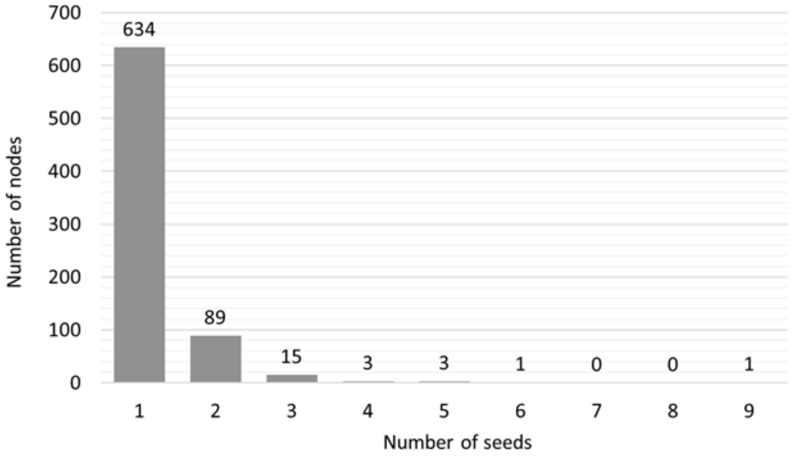


**Supplementary Figure 2. Node degree distribution of the proteins of the global HSP-PPIN based on their connectivity with seeds**
The nodes of the global HSP-PPIN were analysed to calculate the number of seeds to which they connect. The interactors connected to one seed (n=634) were not included in further analyses.

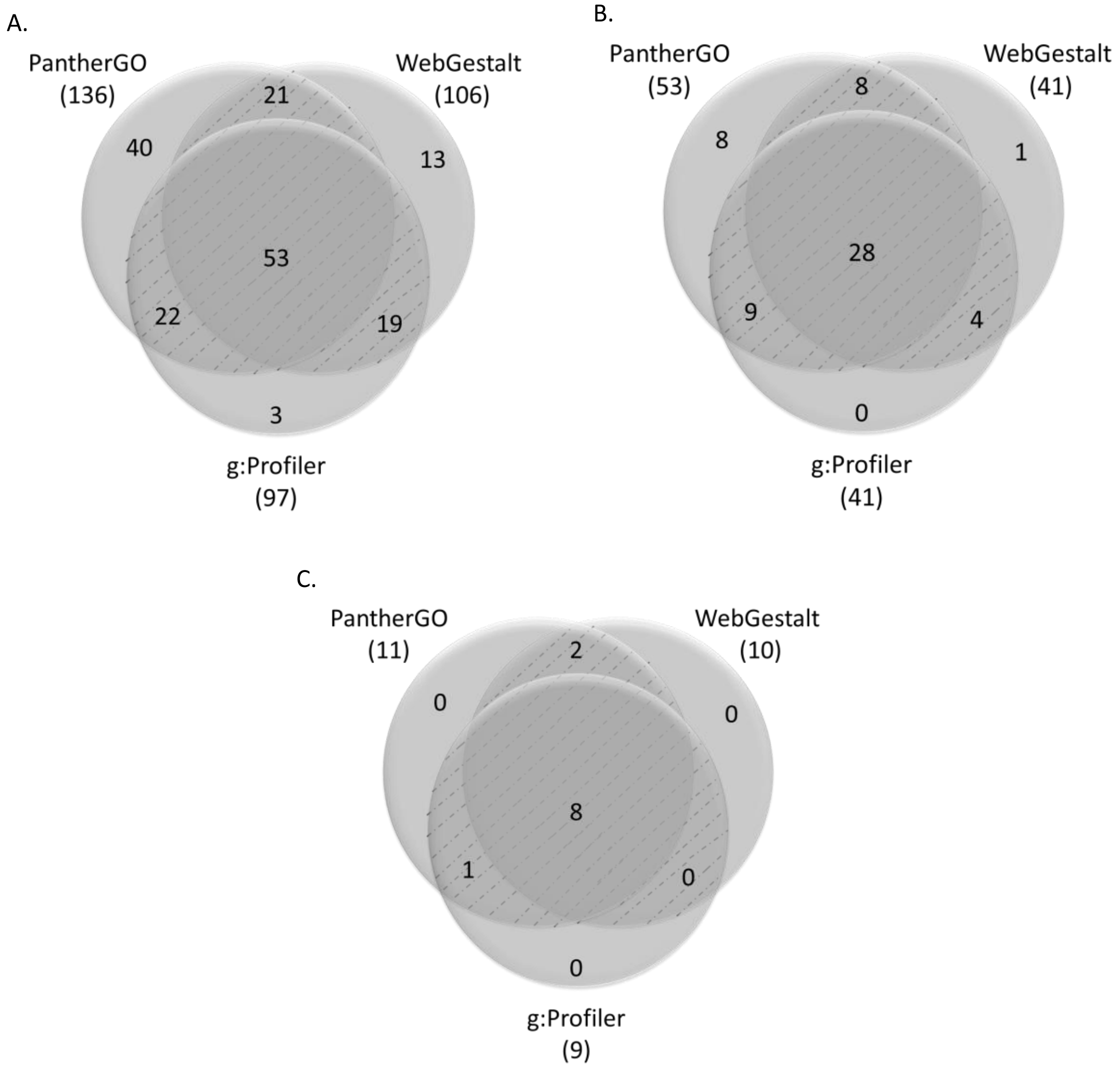


**Supplementary Figure 3. Overlap of the three functional enrichment tools for the analysis of the core HSP-PPIN**
The results from the functional enrichment analysis of the core HSP-PPIN were compared across the functional enrichment tools used, in three levels: single GO-BP terms (n=171) (**A**), semantic classes (n=58) (**B**), and functional blocks (n=11) (**C**).

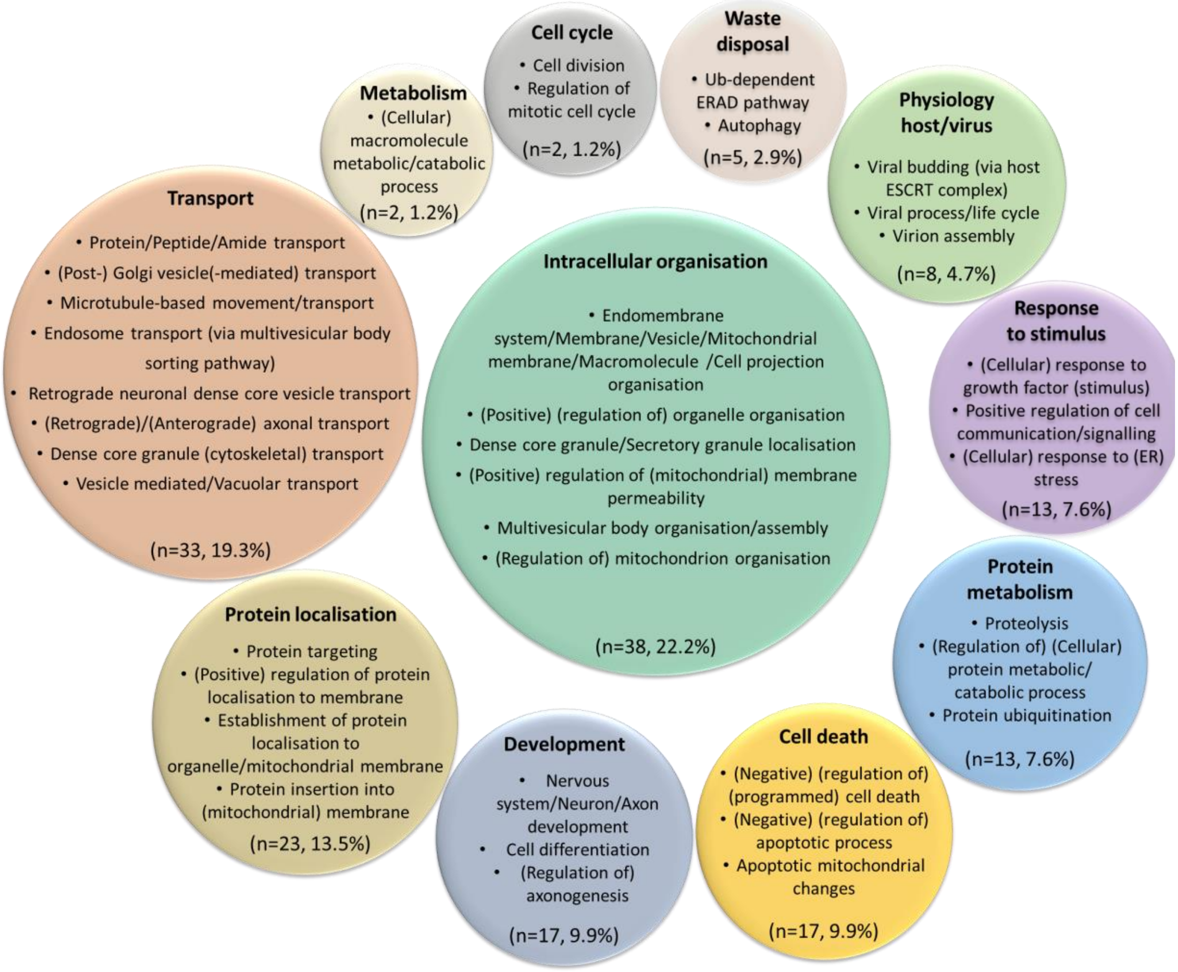


**Supplementary Figure 4. Detailed graphical representation of the functional enrichment of the core HSP-PPIN**
Functional enrichment was performed for the components of the core HSP-PPIN. The resulted GO-BP terms (n=171) (see Supplementary File 3) were grouped into semantic classes and then into functional blocks (name of each circle, bolded). The number and percentage of terms in each functional block was calculated by merging the data from g:Profiler, WebGestalt, and PantherGO as described in Materials and Methods. Examples of GO-BP terms are included inside the circle of each functional block.

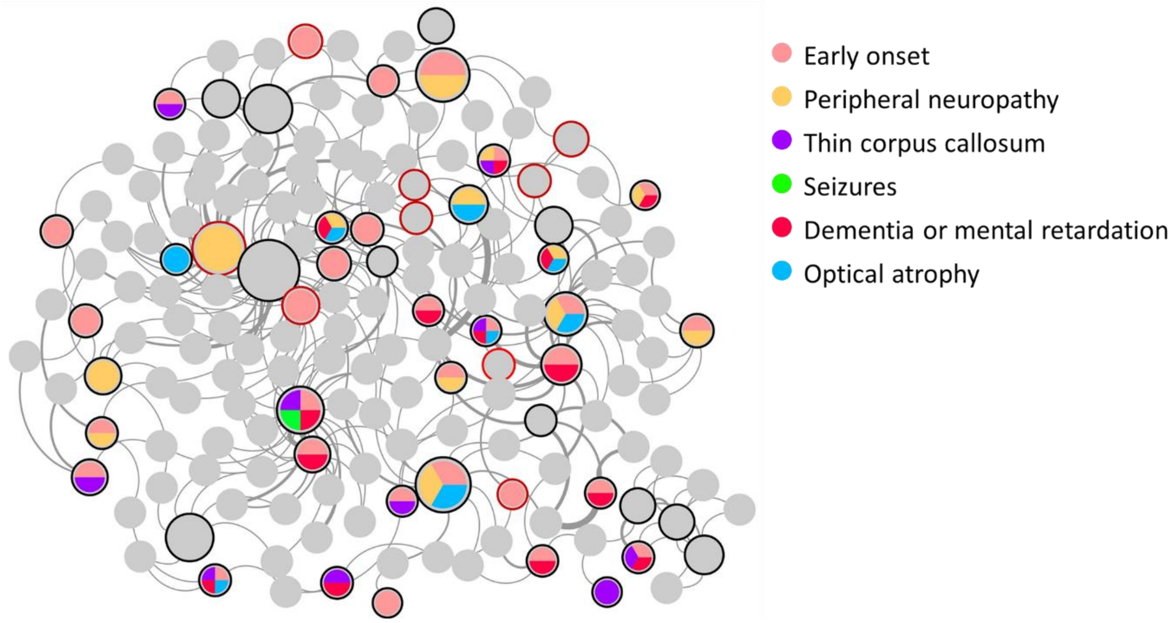


**Supplementary Figure 5. Visualisation of clinical characteristics of HSPs caused by each HSP gene of the core HSP-PPIN**
The presence of clinical characteristics in HSPs is visualised in the core HSP-PPIN through the colour of each node. The colour-function correspondence is located on the top right of the image, while the grey nodes did not present any of the clinical features. The nodes corresponding to the HSP seeds have a black border, while the test seeds have a red border. The size of each node correlates with its degree. The thickness of each edge correlates with its final score as calculated by PINOT.

A.

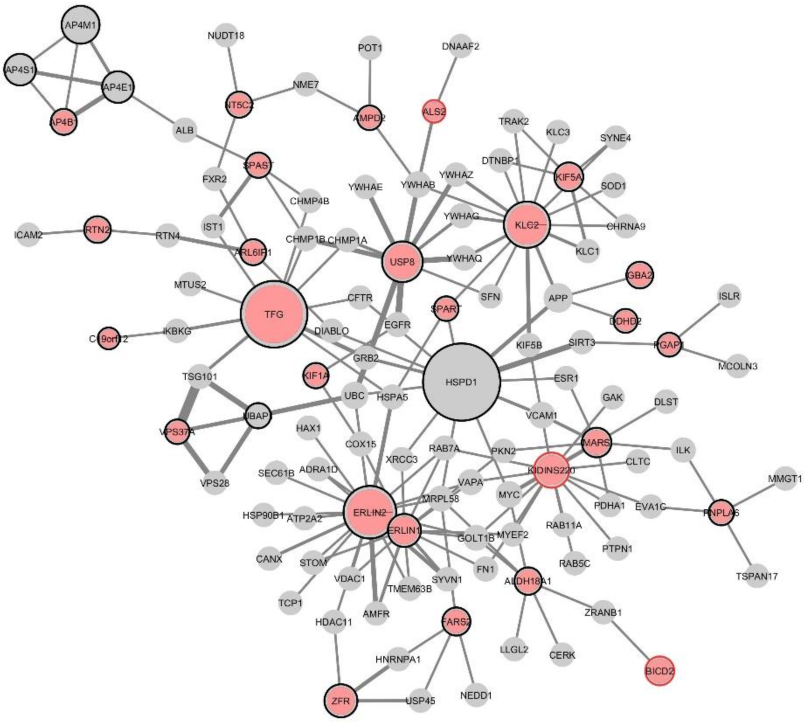

B.

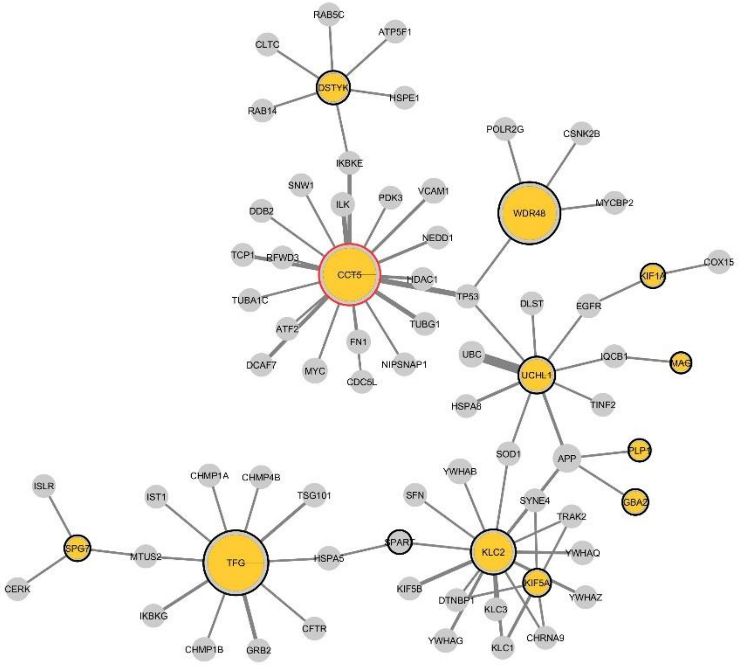
RAB5C
ATP5F1
CLTC
DSTYK
HSPE1
RAB14
IKBKE
POLR2G
CSNK2B
SNW1
ILK
PDK3
VCAM1
DDB2
WDR48
MYCBP2
NEDD1
TCP1
RFWD3
CCT5
HDAC1
TP53
KIF1A
COX15
TUBA1C
DLST
EGFR
ATF2
TUBG1
DCAF7
FN1
MYC
NIPSNAP1
CDC5L
UBC
UCHL1
IQCB1
MAG
HSPA8
TINF2
SOD1
APP
PLP1
ISLR
CHMP1A
CHMP4B
YWHAB
IST1
TSG101
SFN
SYNE4
GBA2
TRAK2
SPG7
MTUS2
TFG
HSPA5
SPART
KLC2
YWHAQ
CERK
KIF5B
KIF5A
YWHAZ
IKBKG
DTNBP1
KLC3
CFTR
YWHAG
CHRNA9
CHMP1B
GRB2
KLC1

C.

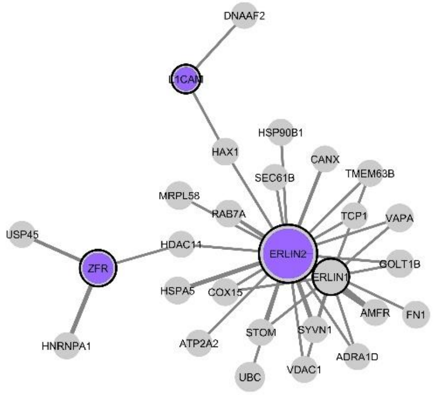
DNAAF2
L1CAM
HSP90B1
HAX1
CANX
SEC61B
TMEM63B
MRPL58
TCP1
RAB7A
VAPA
USP45
HDAC11
ERLIN2
GOLT1B
ZFR
ERLIN1
HSPA5
COX15
AMFR
FN1
SYVN1
STOM
ATP2A2
ADRA1D
HNRNPA1
UBC
VDAC1

D.

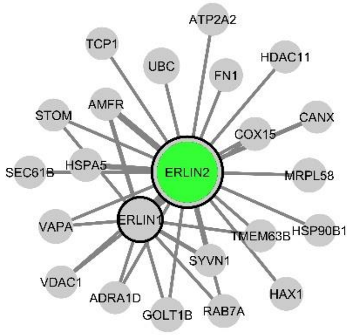
ATP2A2
TCP1
UBC
FN1
HDAC11
AMFR
STOM
CANX
COX15
SEC61B
HSPA5
ERLIN2
MRPL58
VAPA
ERLIN1
TMEM63B
HSP90B1
SYVN1
VDAC1
HAX1
ADRA1D
GOLT1B
RAB7A

E.

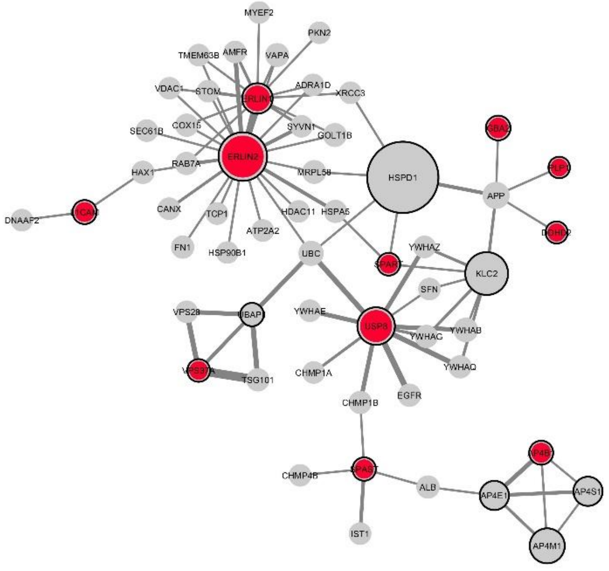
MYEF2
PKN2
TMEM63B
AMFR
VAPA
VDAC1
STOM
ADRA1D
XRCC3
COX15
SYVN1
GOLT1B
SEC61B
RAB7A
ERLIN2
MRPL58
HSPD1
HAX1
APP
DNAAF2
CANX
TCP1
HDAC11
HSPA5
ATP2A2
FN1
HSP90B1
UBC
YWHAZ
KLC2
SFN
VPS28
YWHAE
YWHAB
YWHAG
YWHAQ
TSG101
CHMP1A
EGFR
CHMP1B
CHMP4B
ALB
AP4E1
AP4S1
IST1
AP4M1

F.

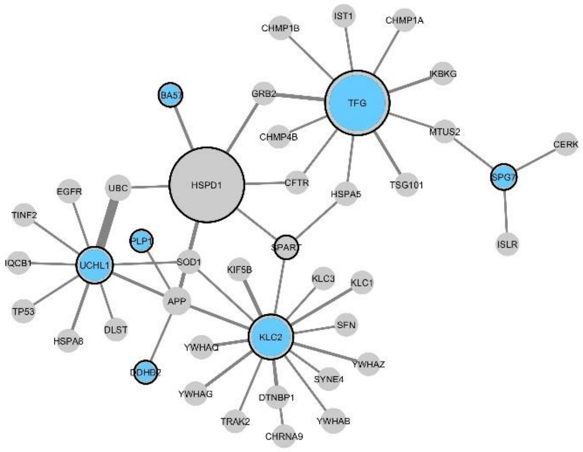


**Supplementaty Figure 6. Comparison of the core network of seeds with the analysed clinical features**
The presence of clinical characteristics in HSPs is visualised in the core networks by the colour of each node for early onset (**A**), peripheral neuropathy (**B**), thin corpus callosum (**C**), seizures, (**D**), dementia or mental retardation (**E**), and optic atrophy (**F**), while the nodes without that clinical feature are grey. The nodes corresponding to the HSP seeds have a black border, while the test seeds have a red border. The size of each node correlates with its degree. The thickness of each edge correlates with its final score as calculated by PINOT.

A.

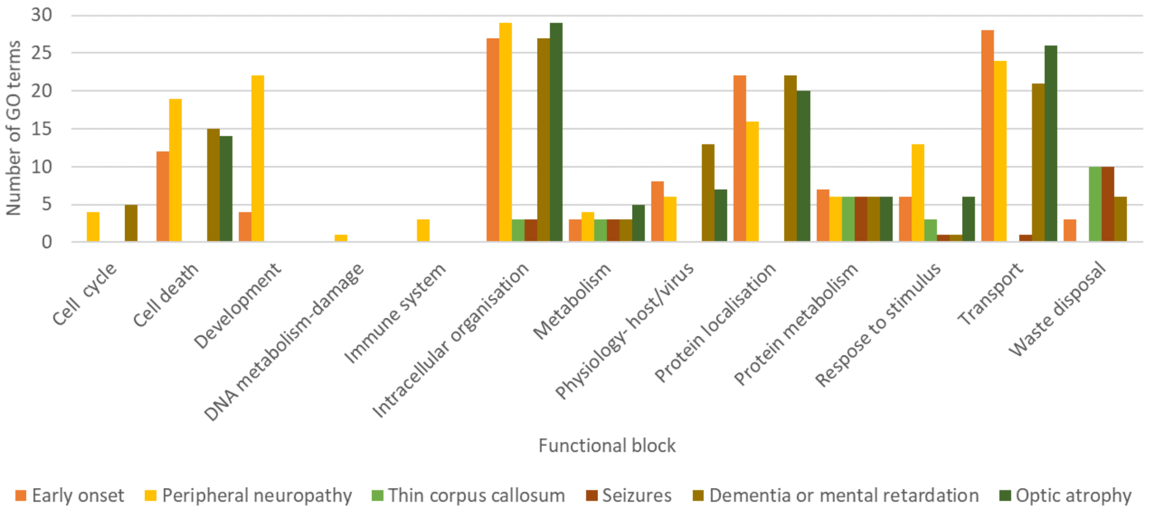


B.

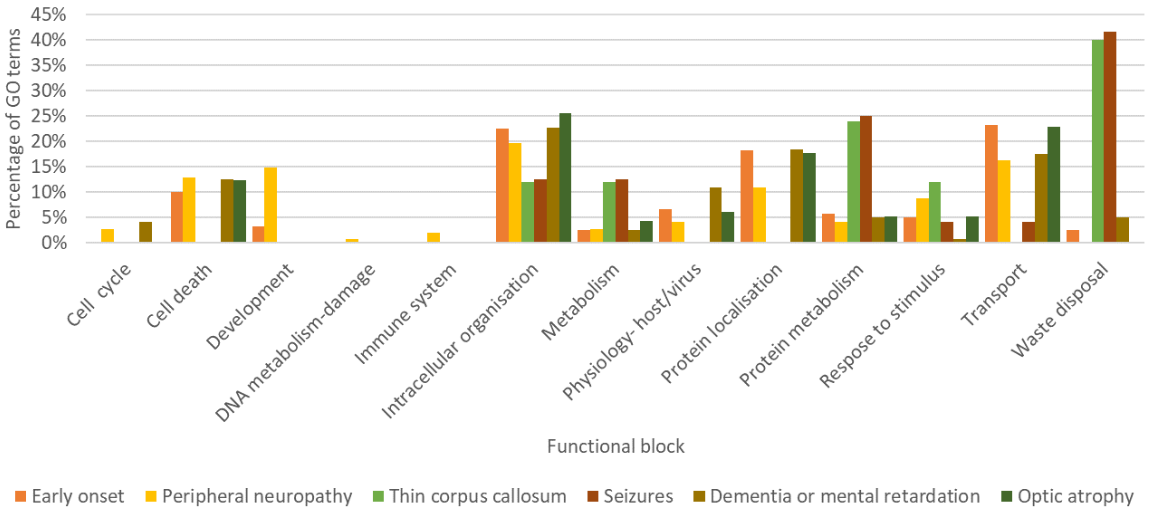


**Supplementary Figure 7. Functional enrichment of the core HSP-PPIN for each group related to a clinical characteristic** The number (**A**) and percentage (**B**) of the GO-BP terms in each functional block is shown for all six groups of genes related to different clinical features. The results are calculated from the functional enrichment data of g:Profiler, PantherGO, and WebGestalt (raw data in Supplementary File 4).

A.

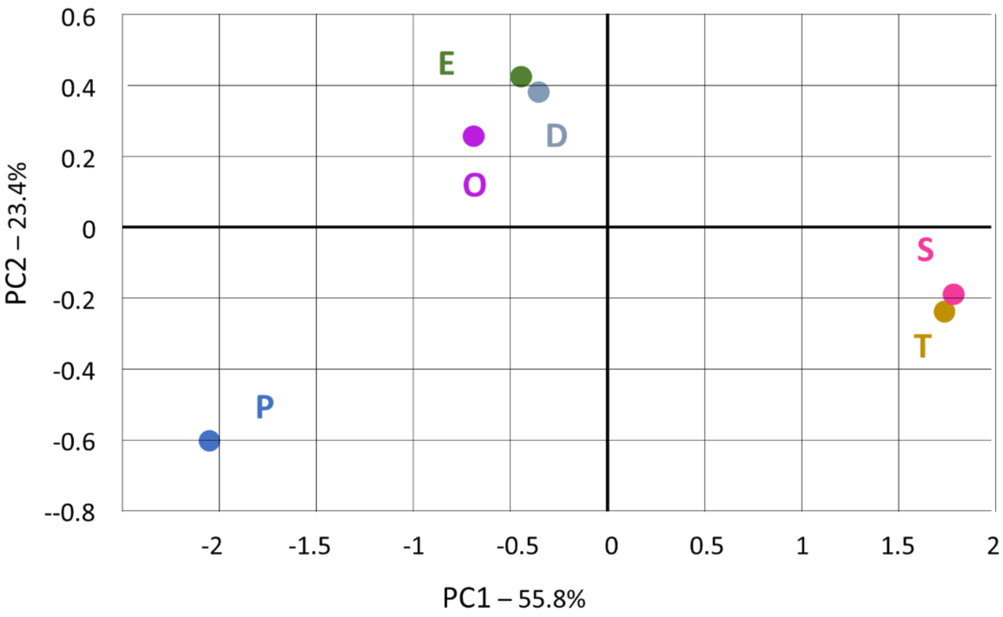
0.6
0.4
0.2
0
-0.2
-0.4
-0.6
--0.8
PC2 – 23.4%
E
D
O
S
T
P
-2
-1.5
-1
-0.5
0
0.5
1
1.5
2
PC1 – 55.8%

B.

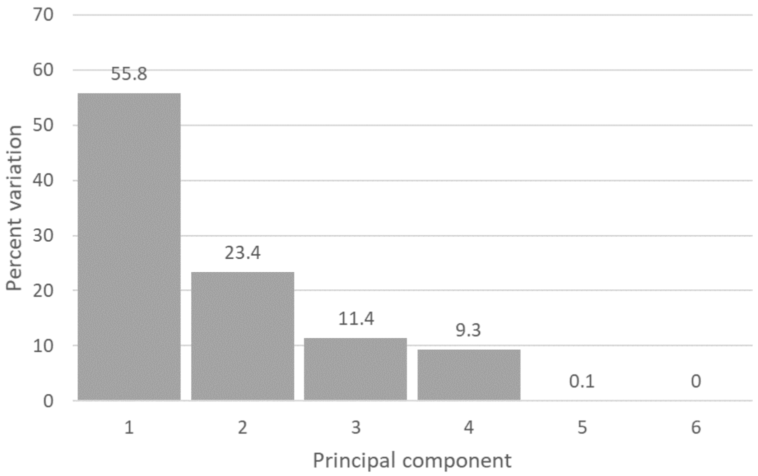
70
60
50
40
30
20
10
0
Percent variation
55.8
23.4
11.4
9.3
0.1
0
1
2
3
4
5
6
Principal component

C.

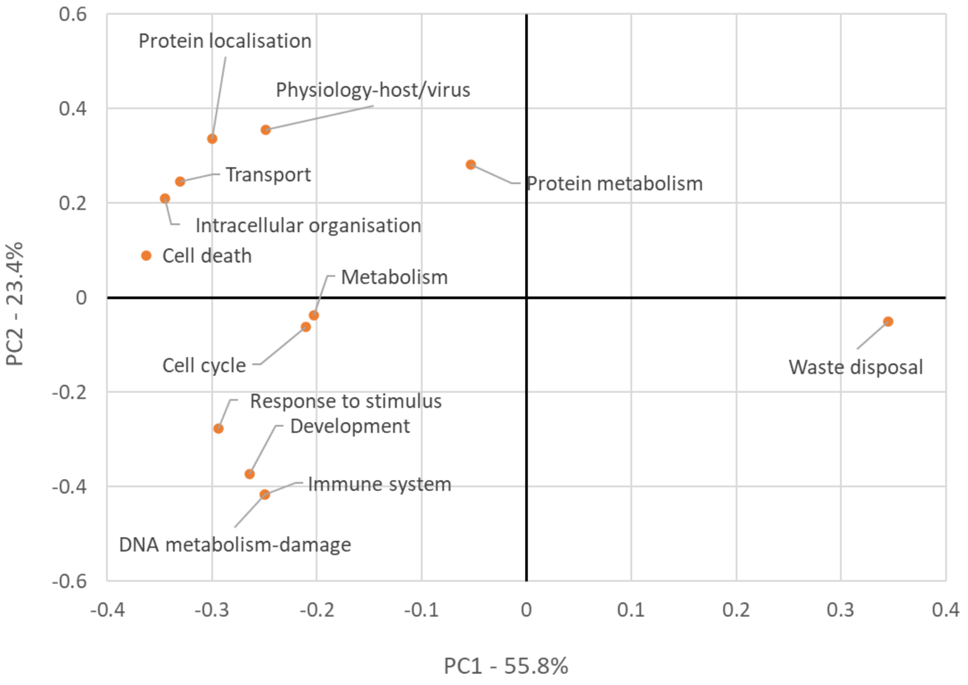
Protein localisation
Physiology-host/virus
Transport
Protein metabolism
Intracellular organisation
Cell death
Metabolism
Cell cycle
Waste disposal
Response to stimulus
Development
Immune system
DNA metabolism-damage
0.6
0.4
0.2
0
-0.2
-0.4
-0.6
-0.4
-0.3
-0.2
-0.1
0
0.1
0.2
0.3
0.4
PC2 - 23.4%
PC1 - 55.8%

D.

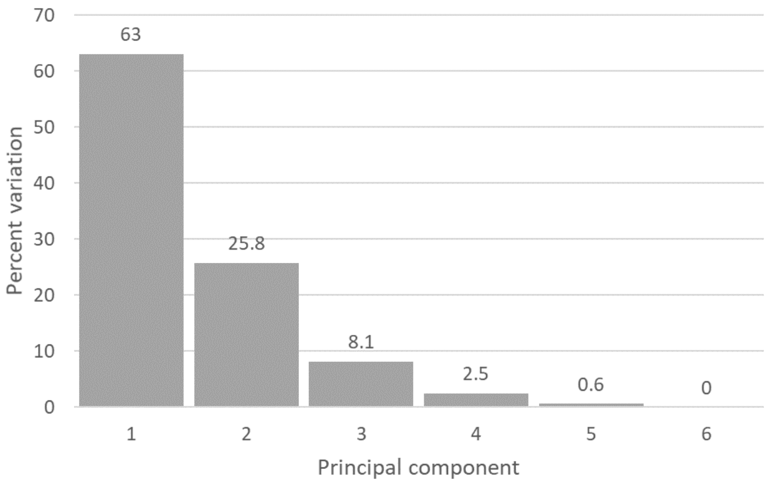
70
60
50
40
30
20
10
0
63
25.8
8.1
2.5
0.6
0
1
2
3
4
5
6
Percent variation
Principal component

E.

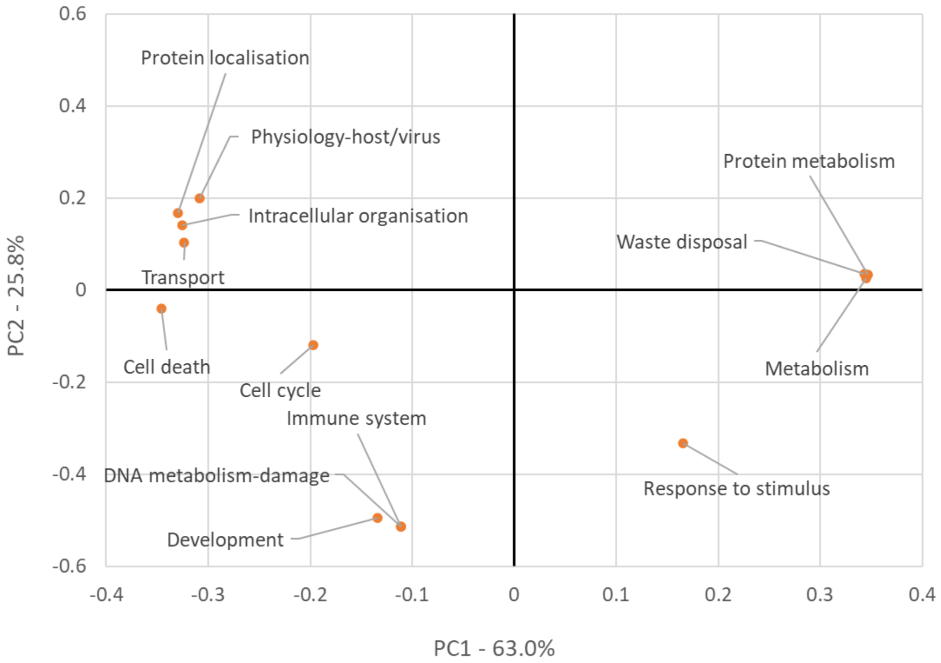


**Supplementary Figure 8. Comparison of the functional profiles of the 6 subdivisions of the core HSP-PPIN based on clinical features using Principal Component Analysis**
The number (**A**, **B**, **C**) and percentage (**D**, **E**) of GO-BP terms for each functional block were analysed with PCA through R. (**A**) The PCA graph is showing the distribution of the gene groups in the PC1 and PC2 axes. The axes have been adjusted based on their respective explained variation. (**B**, **D**) The scree plot is showing the explained variation of the data for PC1 to PC6. (**C**, **E**) The loading scores of each variable (here functional blocks) are plotted against PC1 and PC2, indicating which functions drive the localisation of the gene groups in the PCA graph. E: Early onset, P: Peripheral neuropathy, T: Thin corpus callosum, S: Seizures, D: Dementia or mental retardation, O: Optic atrophy

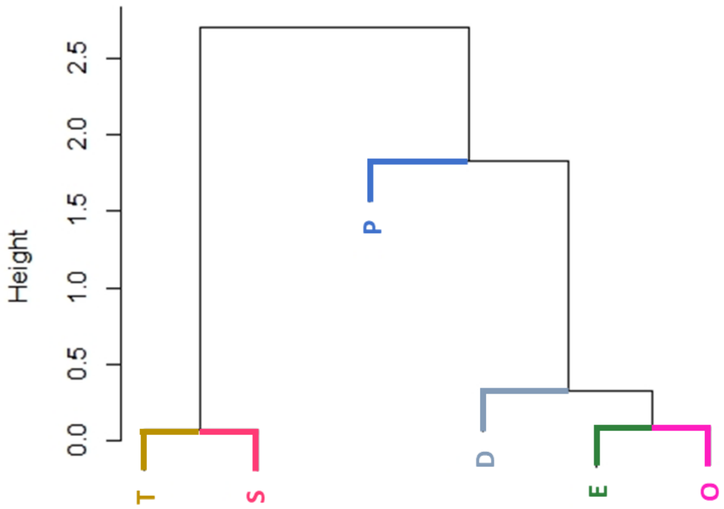


**Supplementary Figure 9. Cluster dendrogram for the number of GO-BP terms in the enrichment of the clinical subnetworks following PCA**
Cluster dendrogram produced based on hierarchical clustering of the gene groups as analysed in Fig 4. E: Early onset, P: Peripheral neuropathy, T: Thin corpus callosum, S: Seizures, D: Dementia or mental retardation, O: Optic atrophy

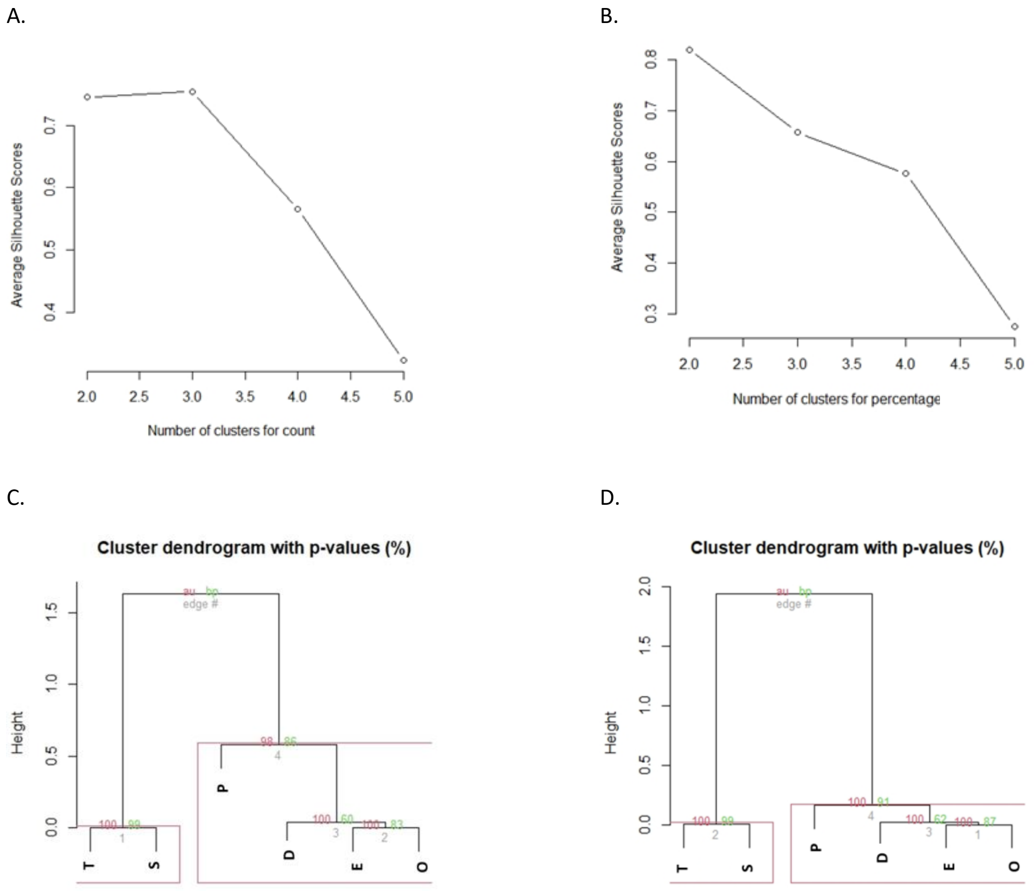


**Supplementary Figure 10. Evaluating the optimal number of clusters using the Silhouette method and the Multiscale bootstrap resampling through R**

The analysis was based on the number (**A**, **C**) and percentage (**B**, **D**) of the GO-BP terms in functional block. The graphs based on the Silhouette method (**A**, **B**) show the optimal number of clusters being the one with the highest value, whereas based on multiscale bootstrap resampling (**C**, **D**) the recommended clusters are framed in red boxes. E: Early onset, P: Peripheral neuropathy, T: Thin corpus callosum, S: Seizures, D: Dementia or mental retardation, O: Optic atrophy

**Supplementary Table 3. Overlap of protein composition within the TS cluster**

| | T | S |
|---|---|---|
| T | | 100.0% |
| S | 82.1% | |

**Supplementary Table 4. Overlap of protein composition within the EPOD cluster**

| | E | P | D | O |
|---|---|---|---|---|
| E | | 60.3% | 96.5% | 81.0% |
| P | 38.6% | | 36.8% | 92.9% |
| D | 48.2% | 28.8% | | 45.2% |
| O | 29.8% | 53.4% | 33.3% | |

| Supplementary Table 5. Overlap of protein composition between the TS and the EPOD cluster | | | | | | |
|---|---|---|---|---|---|---|
| | E | P | T | S | D | O |
| E | | | 96.4% | 100.0% | | |
| P | | | 17.9% | 21.7% | | |
| T | 23.7% | 6.8% | | | 43.9% | 4.8% |
| S | 20.2% | 6.8% | | | 40.4% | 4.8% |
| D | | | 89.3% | 100.0% | | |
| O | | | 7.1% | 8.7% | | |

A.

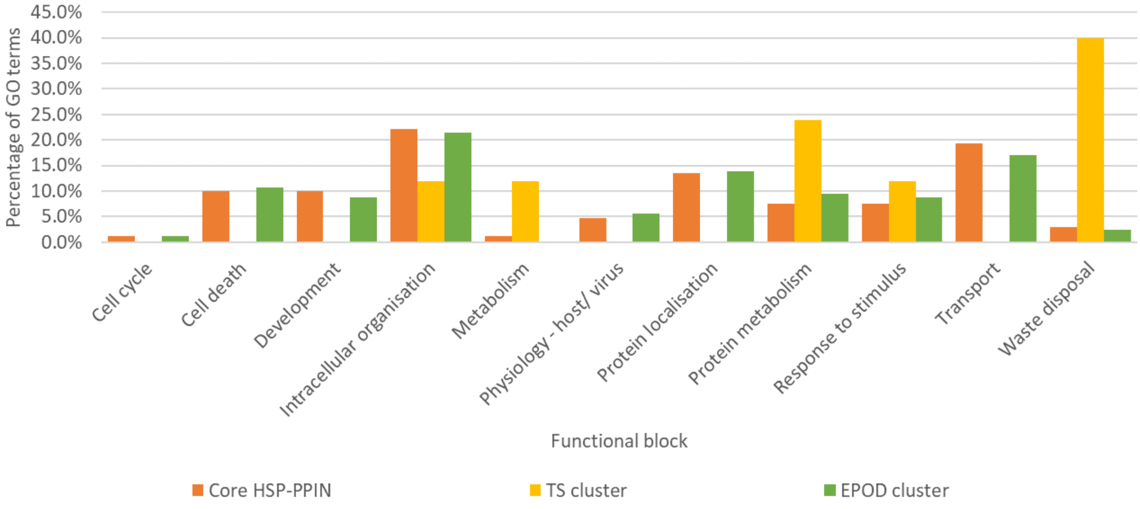


B.

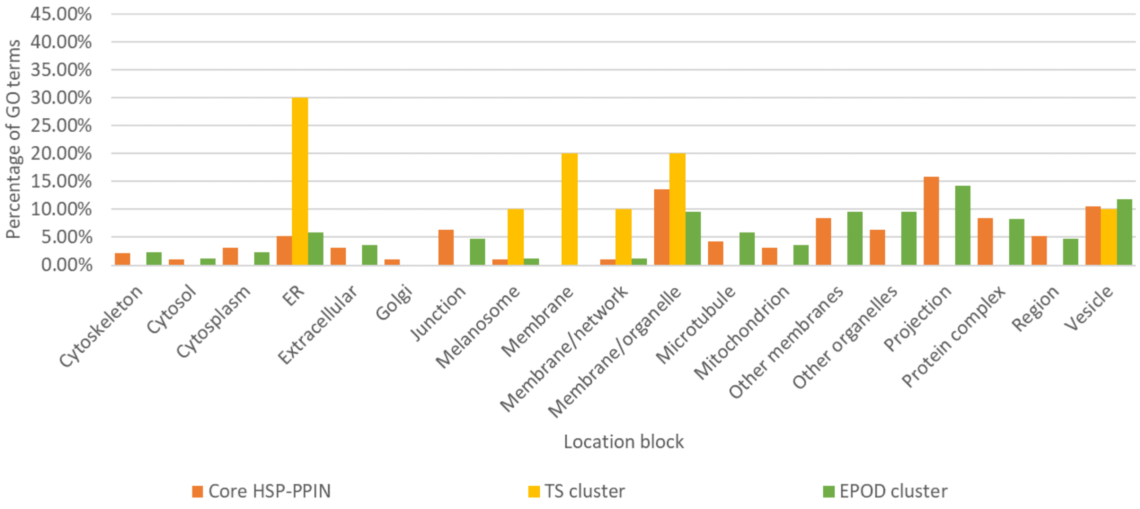

C.

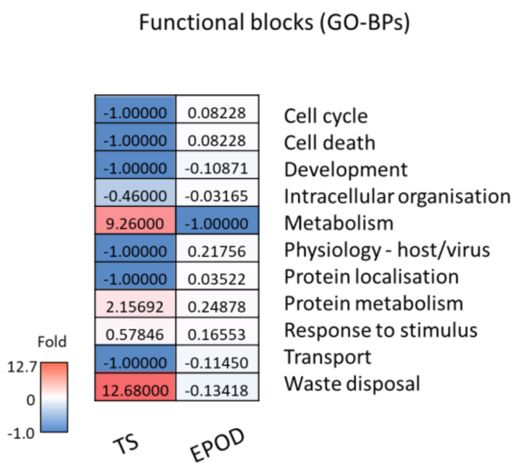


D.

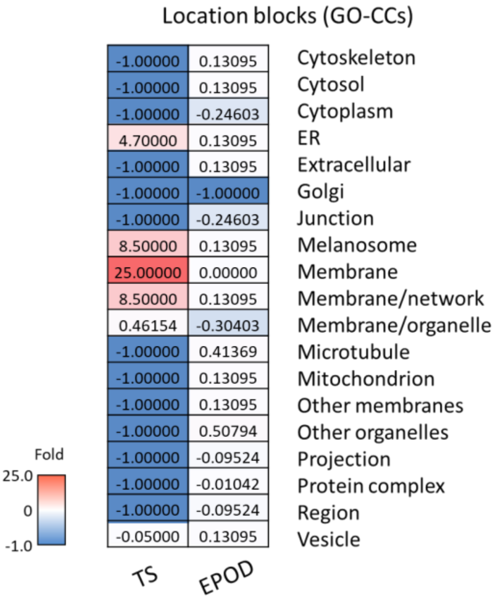


**Supplementary Figure 11. Comparison of the enrichment profile of the two clusters of the core HSP-PPIN**
The analysis is based on the percentage of GO terms in each functional or location block as resulted from the enrichment of GO-BPs (**A**, **C**) and GO-CCs (**B**, **D**). The comparison is expressed as a fold change compared to the profile of the core HSP-PPIN (**C**, **D**).
